# Population codes for biological stereopsis extend beyond correlation-based binocular disparity computations

**DOI:** 10.64898/2026.04.21.719805

**Authors:** Bayu Gautama Wundari, Ichiro Fujita, Hiroshi Ban

## Abstract

Binocular stereopsis depends on comparing the images seen by the two eyes. Although correlation-based models explain responses of individual binocular neurons in primary visual cortex (V1), it remains elusive whether population representations formed by local correlation activity patterns can support depth perception under ambiguous inputs. Using psychophysics, fMRI, and neural network modeling, we tested human stereopsis with dynamic anticorrelated stimuli that were dominated by binocular mismatches. Humans reliably perceived reversed depth as predicted by correlation-based computations, yet population representations consistent with this percept were detected in V3A, not V1. Shallow and deep neural networks constrained by correlation-like binocular interactions did not capture the full pattern of human depth judgments. Analyses of their internal representations showed greater representational overlap, whereas deep architectures not constrained to explicit correlation interactions exhibited less entangled representations and better aligned with human behavior. These findings suggest that biological stereopsis may rely on population coding beyond correlation-like computations.

**Significance Statement:** The brain must infer depth from binocular inputs that are inherently ambiguous. Although correlation-based models explain disparity tuning of individual neurons in primary visual cortex (V1), whether these local mechanisms support perceptual inference at the population level remains unclear. Using psychophysics, fMRI, and neural network modeling, we show that population representations consistent with perceived depth under ambiguity were detected in mid-dorsal area V3A, not V1. Analyses of how neural networks encode multiple features showed that correlation-based computations represent features into overlapping activity patterns that may constrain downstream readout and degrade depth estimates. In contrast, models not constrained by explicit interocular correlation maintained more distinct population codes and closely matched human perception. These findings suggest that architectures constrained to explicit correlation-like processing can form population representations that are suboptimal to explain human stereopsis, motivating hybrid mechanisms.

## Introduction

Our brain compares two slightly different perspectives of the world (binocular disparity) when seeing with two eyes to construct a unified 3D structure of the viewed scene. To infer depth from disparity, the visual system must form a representational space capable of solving the stereo correspondence problem: identifying which image patches in the two eyes originate from the same point in space (true matches) and which do not (false matches) ^1^. The computation underlying this process is challenging because the visual brain must not only extract binocular information from inputs that are inherently ambiguous and noisy, but also organize them into a representational format that supports robust depth perception.

A major advance in the understanding of biological stereopsis came from correlation-based computations, formalized in the binocular energy model (BEM ^2^) and its extensions ^3–7^. At the core, these correlation theories of stereopsis propose that binocular neurons encode disparity-related signals by correlating image patches between the two eyes, conveying information about true and false disparities ^2–10^. These models have been highly influential in explaining key physiological properties of single V1 neurons, particularly the inversion of disparity tuning when binocular images are anticorrelated (contrast polarity is reversed between the two eyes ^3,11,12^).

However, correlation-based models focus on local nonlinear computations at the level of single neurons. Thus, they provide limited insights into how such local signals are transformed into population codes to extract disparity information from ambiguous inputs. Indeed, much of our knowledge about stereo coding in V1 derives from neurons whose tuning is well described by Gabor functions ^13–16^, a hallmark of the disparity energy model ^2,14^. This evidence is essential for explaining local response properties of binocular neurons, but it remains unclear whether population codes based solely on their local activity patterns can underlie human depth judgments when binocular inputs are ambiguous.

Here, we tested if population representations encoded exclusively by correlation-like computations can support human-like depth judgments under ambiguous inputs. Using random dot stereograms (RDSs, Fig. 1a) to isolate binocular disparity cues, we manipulated the balance between true and false matches, and measured both perceived depth and brain responses using fMRI. This stimulus configuration allowed us to examine how local correlation-like disparity signals can be transformed into population codes that support reliable depth perception. Humans perceived reversed depth for dynamic anticorrelated stimuli, consistent with predictions from correlation-based computations (Fig. 2a). However, multivariate fMRI analysis revealed voxel representations consistent with this percept were linearly decodable in a mid-level visual area V3A, but not in V1 (Fig. 2b, c). In parallel, shallow and deep neural network models that were constrained by correlation-based mechanisms (Fig. 3a) captured aspects, but not the full pattern of human depth judgments under ambiguous binocular inputs (Fig. 4). In contrast, deep architectures that preserved monocular information more closely matched human behavior (Fig. 4).

**Figure 1:**
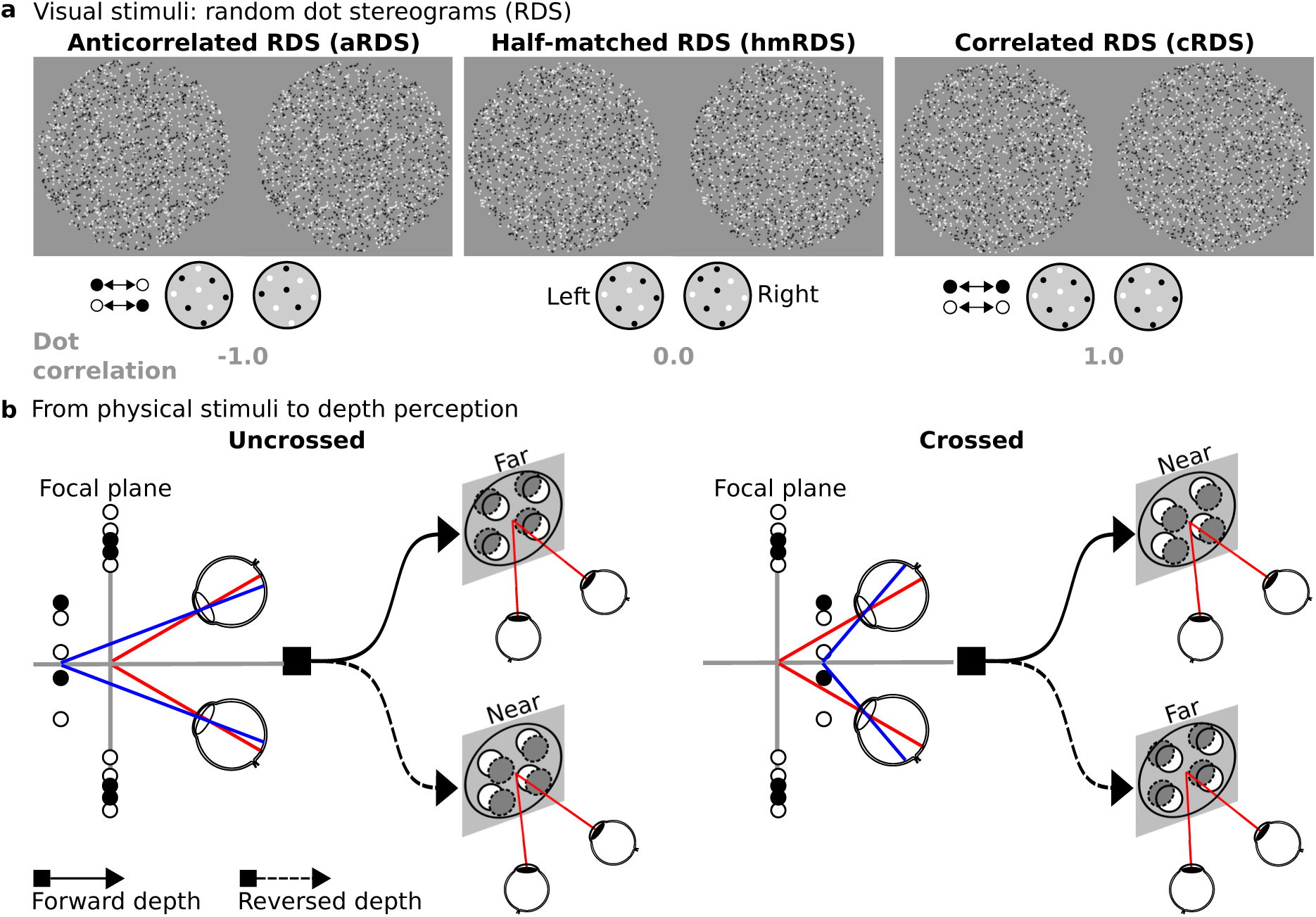
Visual stimuli and perceptual outcomes in RDSs. **a** Schematic of three RDS types: correlated RDSs (cRDS; all dots contrast-matched between the two eyes, binocular dot correlation = +1), half-matched RDSs (hmRDS; 50% contrast-matched and 50% contrast-reversed dots, correlation = 0), and anticorrelated RDSs (aRDS; all dots contrast-reversed between the eyes, correlation = –1). Each stimulus comprises four identical circular RDS apertures arranged in quadrants within the zero-disparity background. **b** Two scenarios of perceiving depth in RDSs. Forward depth corresponds to the veridical depth perception: crossed disparity perceived as near and uncrossed as far (solid black curves). Reversed depth is the opposite percept (dashed black curves). Red lines denote fixation at the focal plane, and blue lines illustrate the projection of dots with crossed or uncrossed disparities onto the retinae.

**Figure 2:**
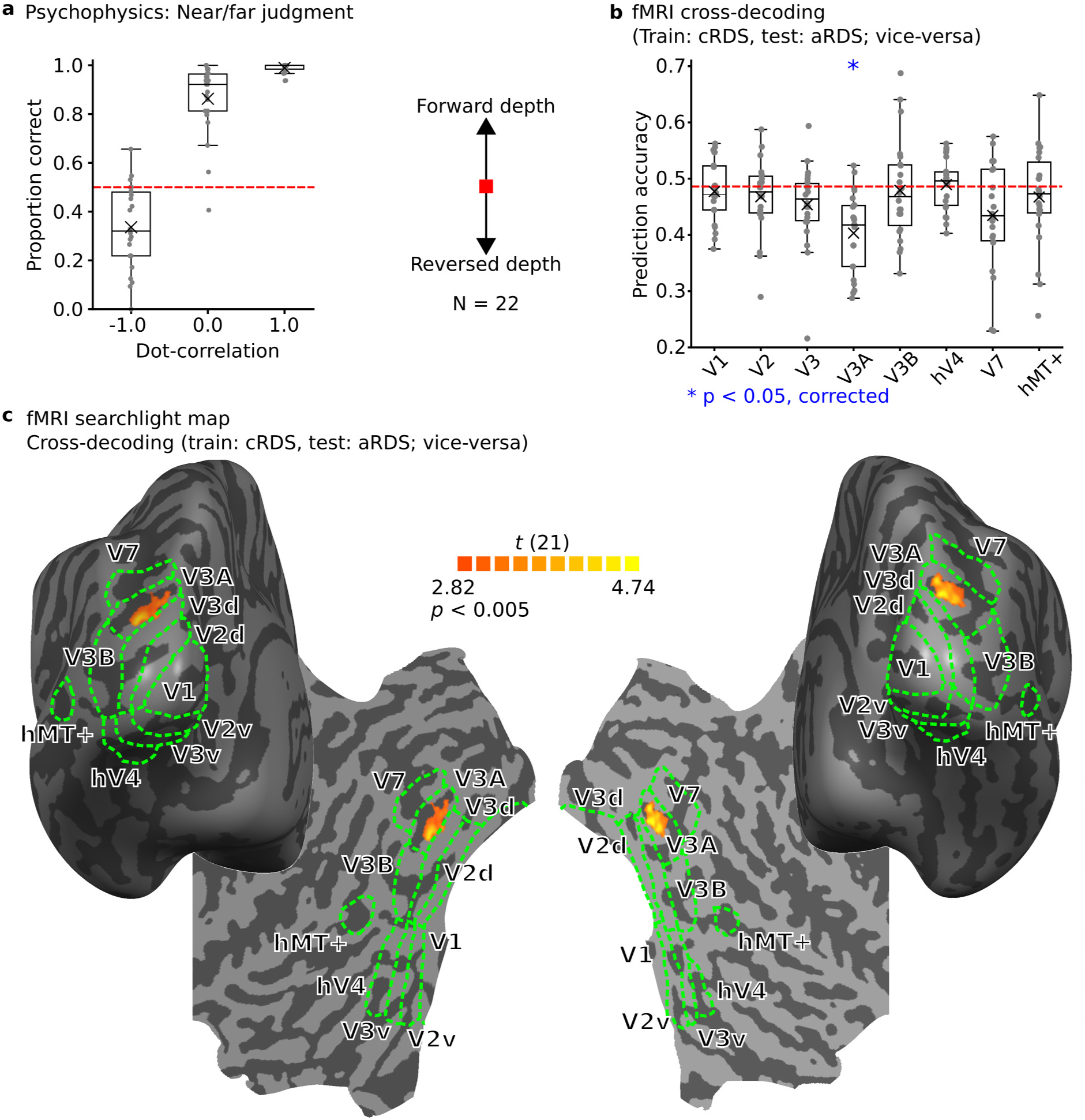
Human depth judgments and cortical representations of reversed depth. **a** Near/far discrimination performance for dynamic aRDS, hmRDS, and cRDS (*N* = 22). Dots represent individual participants; black crosses denote the mean; red dashed lines indicate chance (0.5). The proportion correct below chance denotes reversed depth. **b** Cross-decoding performance (training on cRDS and testing on aRDS, *vice versa*) within each visual area for the same participants. Dots represent individual data points; red dashed lines indicate baseline (permutation-based chance level). Blue asterisks mark significant performance below the baseline (adjusted *p <* 0.003125). **c** Searchlight map on a representative individual’s cortical surface (posterior and flattened views). Sulci are shown in darker gray than gyri; dashed green lines demarcate the retinotopic boundaries. Color coding represents *t*-values from a group-level searchlight analysis for crossdecoding between cRDS and aRDS (6-mm radius searchlight sphere, linear SVM, threshold: *p <* 0.005, uncorrected).

**Figure 3:**
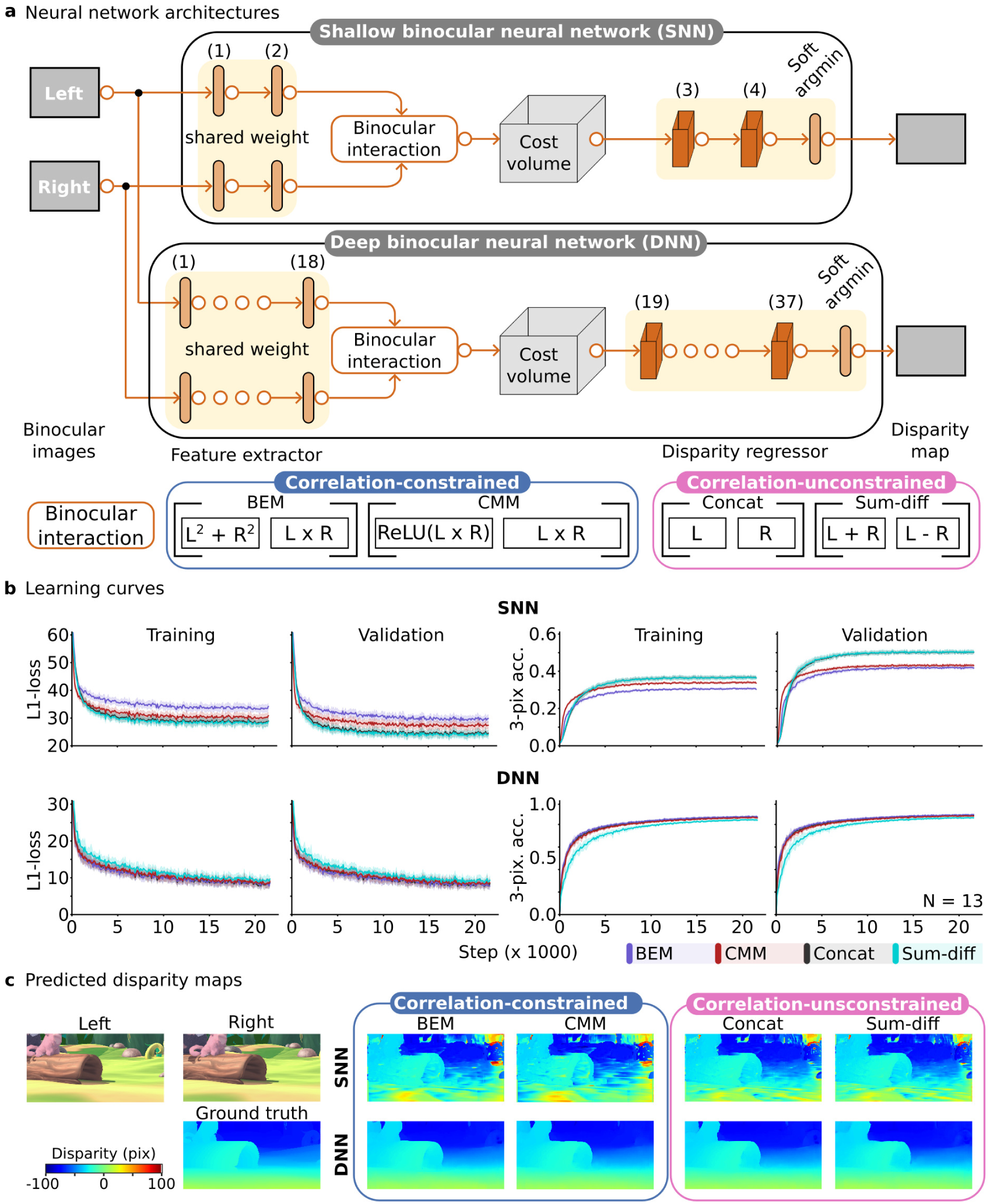
Neural network architectures and performance. **a** Schematic of the shallow neural network (SNN) and deep neural network (DNN). Each model was implemented with one of four binocular interactions: Concat, BEM ^2^, CMM ^5,22^, or Sum-diff ^29,30^. **b** Training and validation learning curves for each binocular interaction, trained on Monkaa datasets for 10 epochs and averaged across 13 random seed numbers. Shaded areas represent one standard deviation across seeds (*N* = 13). The training errors were higher than those of validation due to data augmentation applied during training (see Methods). **c** Example predicted disparity maps from SNNs and DNNs with each binocular interaction, evaluated on the Monkaa validation dataset. Reddish colors represent pixels closer to the camera, bluish colors represent pixels farther away.

**Figure 4:**
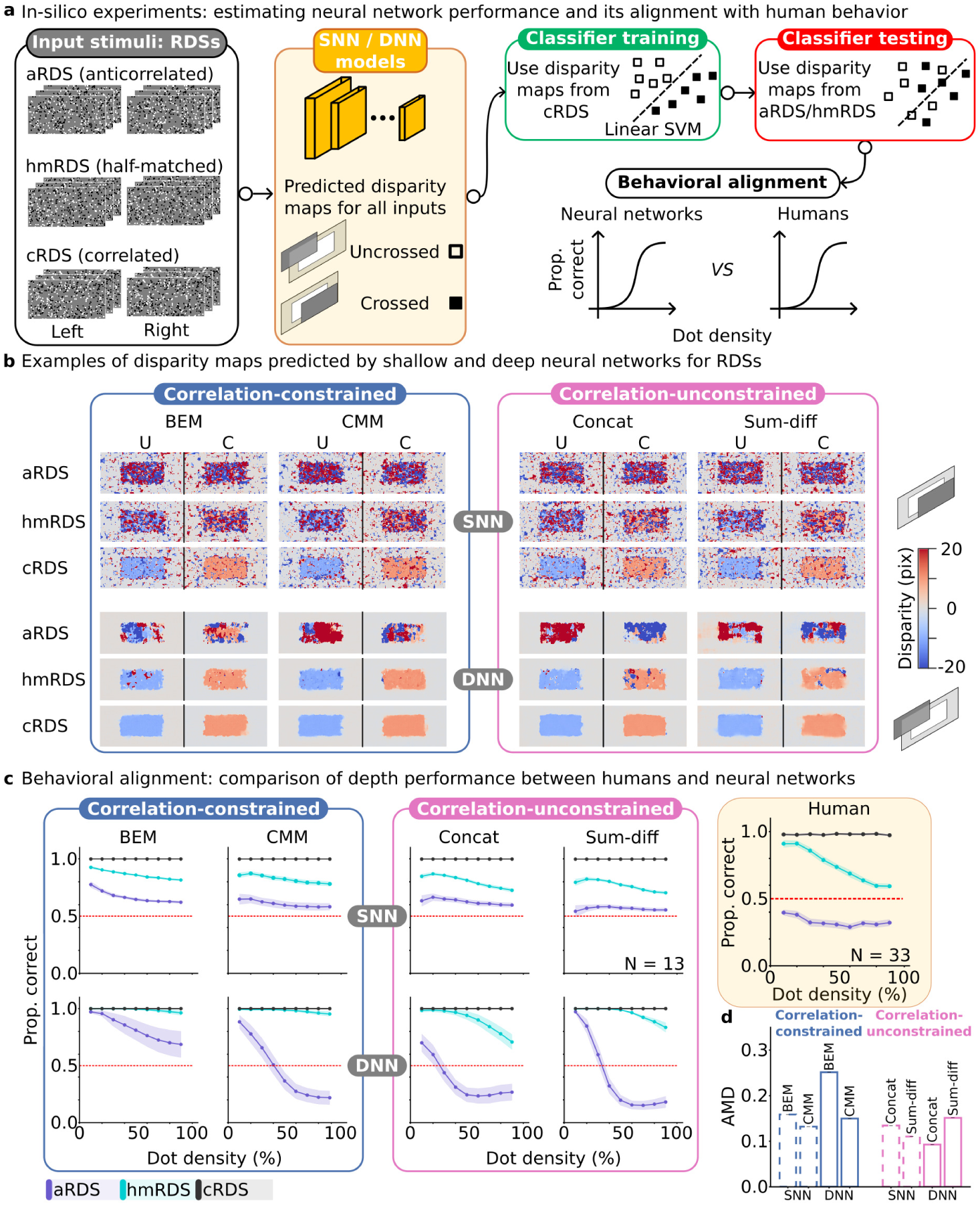
Behavioral alignment between models and humans. **a** Schematic of the *In-silico* evaluation procedure. Each model was presented with RDSs varying in dot correlation (aRDS, hmRDS, and cRDS) and dot density (0.1, 0.2, …, 0.9). Disparity maps predicted for cRDS were used to train a linear SVM to classify crossed *vs.* uncrossed disparities, then tested on disparity maps for aRDS and hmRDS. Classification accuracy served as the model’s metric of depth performance. **b** Representative disparity maps for RDSs predicted by SNNs and DNNs for each binocular interaction, shown for stimuli with target disparity of 10 pixels and dot density of 50%. Reddish colors represent pixels closer to the camera, bluish colors represent pixels farther away. **c** Near/far discrimination performance for neural networks and humans across dot densities for aRDS (purple), hmRDS (cyan), and cRDSs (black). Shaded areas indicate the standard error across participants (humans, *N* = 33) or random seeds (models, *N* = 13). **d** Absolute mean difference (AMD) between human and model psychophysics, averaged across dot densities and dot match levels. Dashed outlines denote SNNs; solid outlines denote DNNs. Blue indicates correlation-based interactions; magenta indicates correlation-unconstrained interactions.

To understand why model behavior differed, we analyzed their internal representations by measuring the degree of representational overlap across layers. We employed representational analysis inspired by superposition theory from AI-interpretability research as a diagnostic tool, not as direct evidence that biological circuits implement the same coding scheme as artificial networks. This theory of representations asks how to pack many task-relevant features in a few dimensions ^17^, making it suitable to gain insight into how correlation signals are organized across layers. The analysis showed stronger representational overlap in models that incorporated explicit correlation-like interactions (Fig. 5), compatible with their poor alignment with human behavior (Fig. 4c, d).

**Figure 5:**
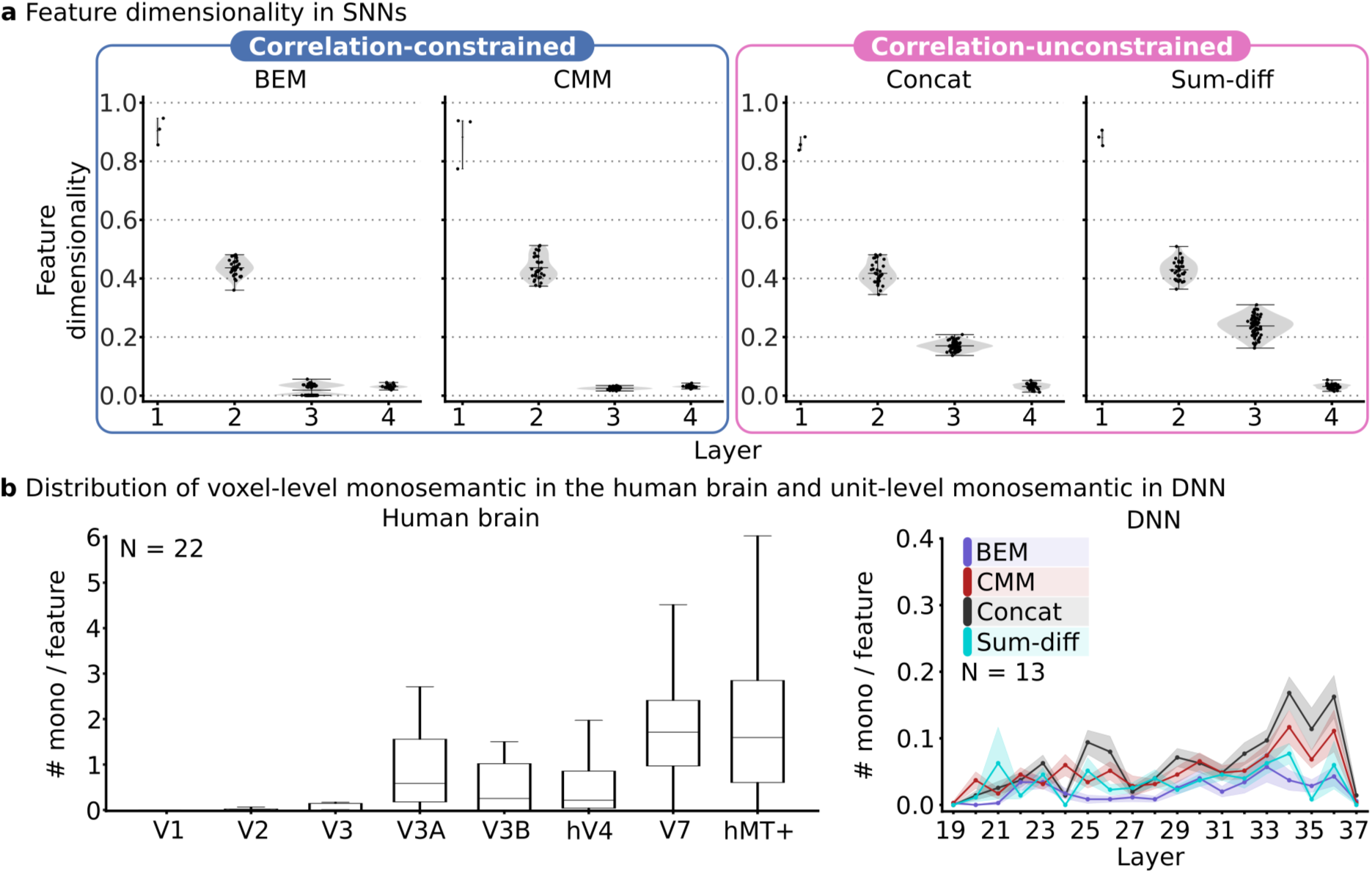
The representational analysis in SNNs, DNNs, and human visual cortex. **a** Feature dimensionality analysis in SNNs, quantifying the fraction of embedding dimensions used to encode distinct features. **b** Distribution of voxel-level monosemantic per feature in the human visual cortex (*N* = 22 participants) and unit-level monosemantic per feature in DNNs (*N* = 13 random seeds). Monosemanticity measures the degree to which units (or voxels) selectively encode a single feature. In both metrics, lower scores suggest greater representational overlap among feature representations.

Together, our study provides a possible computational rationale for why population codes encoded exclusively by correlation-like computations may not fully explain human depth perception. Correlation-based computations capture important local disparity representations in V1, including false matches ^18,19^. However, architectures relying solely on correlation-like interactions exhibited greater representational overlap and demonstrated poor alignment with human behavior, whereas architectures preserving monocular information before later integration better matched human depth judgments. This gives insight into why brain representations consistent with reversed depth perception, a signature of correlation models, were explicit in V3A but not detected at voxel scale in V1. The visual brain may not solve the correspondence problem using a single correlation-like code, but by preserving and transforming multiple binocular formats across areas.

## Results

Participants viewed dynamic RDSs with three levels of binocular correlation: anticorrelated (aRDSs), half-matched (hmRDSs), and correlated (cRDSs), while we measured their behavioral and brain responses (Fig. 1a). We expected two scenarios of perceptual depth from RDSs: forward and reversed (Fig. 1b). Participants experienced forward depth when fixating at the focal planes (Fig. 1b, red lines), points at uncrossed disparities project inwardly and are perceived as far, whereas points at crossed disparities project outwardly and are perceived as near (Fig. 1b, solid black curves). Conversely, reversed depth occurs when uncrossed disparities elicit near perception and crossed disparities elicit far (Fig. 1b, dashed black curves). We performed multivariate analysis to identify the brain region whose population representations were consistent with reversed depth. We then modeled disparity processing using shallow and deep neural networks (SNNs and DNNs, respectively) to examine how different binocular interactions affected model predictions and their alignment with human depth judgments.

### Participants experienced reversed depth when viewing dynamic aRDSs

Participants discriminated crossed disparity as near and uncrossed as far almost perfectly for cRDSs (mean accuracy = 0.989 ± 0.004 SE), well above chance for hmRDSs (mean accuracy = 0.863 ± 0.032 SE), but fell below chance for aRDSs (mean accuracy = 0.336 ± 0.037 SE; Fig. 2a). The below-chance performance for aRDS stimuli indicates participants perceived depth in reverse. However, reversed depth was not observed with static aRDSs (see Fig. S1 and previous studies ^20–22^), suggesting the role of temporal dynamics in inferring disparity. These results suggest that, under dynamic viewing, false match signals were not fully discarded ^4,6,10,11,16,18,23^, rather, could influence downstream neurons for depth inference.

### Reversed depth representations were reliably detected in a mid-dorsal region V3A

Neural signature of reversed depth is characterized by the inversion of disparity tuning between cRDSs and aRDSs. Such inverted tuning at the single-neuron level has been predicted by the correlation-based models (BEM and its extensions ^2,3,5,7^) and observed in V1^2,11^ and some other areas ^24–26^. To identify which cortical areas in the human brain exhibit this inversion at the population level, we applied cross-decoding analyses to our fMRI data ^22^. A linear classifier (support vector machine, SVM) was trained on multivoxel BOLD response patterns elicited by cRDS to classify crossed *vs.* uncrossed disparities and tested on aRDS-induced response patterns, and (*vice versa*). We performed this analysis in retinotopically defined regions of interest (ROIs) V1, V2, V3, V3A, hV4, hMT+, V7^22^.

Only V3A exhibited cross-decoding accuracy significantly below the baseline (permutation-based chance performance) (mean accuracy = 0.403, *p <* 0.001; Fig. 2b), indicating an inverted relationship at the population level between activity patterns evoked by cRDSs and aRDSs. Interestingly, cross-decoding in V1 did not differ from the chance (mean accuracy = 0.48, *p* = 0.22). A whole-brain searchlight analysis ^27^ for cross-decoding between cRDS and aRDS brain response patterns (6 mm radius sphere, a linear SVM as the classification algorithm) only detected significant clusters in V3A and not outside our retinotopically defined ROIs (Fig. 2c), confirming our ROI-based analysis (Fig. 2b).

Together, these results suggest two points. First, population-level disparity representations in V3A are consistent with reversed depth perception. Second, although the inverted relationship has been observed at the single-neuron level in V1^11,12^ and can be explained by correlation-based models ^3,7,13^, it could not be explicitly decoded at the population level by our linear classifier in the same area.

### Examining the correlation mechanisms in shallow and deep neural networks

Inverted disparity tuning observed in single V1 neurons ^2,11^ signifies a hallmark of binocular correlation models. However, our preceding multivariate analysis showed that this signature was undetected at the population level in V1 (Fig. 2b, c). To investigate this discrepancy, we extended correlation mechanisms by incorporating them into SNNs and DNNs. We then evaluated how they orchestrate the representations at the population level and compared their performance with human depth judgments.

Neural network modeling provides a data-driven approach that avoids feature engineering based on predefined receptive field (RF) structures (e.g., Gabor filters) which are assumed in the BEM ^2^ and its variants ^3,5,7,19^. Although the Gabor RFs are well supported by neurophysiological findings in explaining the responses of a subset of V1 neurons, such a specific prior assumption may limit neurons to encode disparity efficiently in a certain kind of environment ^28^, and may not generalize well under complex and ambiguous viewing conditions. It is therefore unlikely that fixed filter assumptions can generate population codes capable of extracting all stereo information in natural scenes. Data-driven neural networks allow us to test this computational constraint without relying on any prior assumption about RFs as the networks learn the filter weights from the stereo data, while still retaining architectural and training priors that we explicitly control across model variants.

To approximate correlation-based binocular processing within the networks, we incorporated binocular interactions analogous to those used in correlation models (Fig. 3a). Concretely, network models with correlation binocular interactions included: the BEM ^2^ which combined the monocular energy terms with the interocular products [*L*^2^ + *R*^2^*, L* × *R*], and the correlation-and-matching model (CMM ^5,22^) which combined the interocular terms with their rectified counterparts [*L* × *R, ReLU* (*L* × *R*)]. The *L* and *R* denote the convolutional layer outputs from the left and right channels, respectively.

As comparison architectures, we implemented correlation-unconstrained binocular interactions that did not explicitly compute interocular correlation. These correlation-unconstrained architectures included: the Concat architecture which concatenated the left and right convolutional layer outputs [*L, R*], and the Sum-diff architecture (summing-and-differencing channels model ^29,30^) which concatenated summation and difference channels [*L* + *R, L* − *R*]. These architectures preserved binocular information without explicitly computing cross-correlation-like products. All training hyper-parameters were matched, except for the binocular interaction types and the number of layers in SNNs and DNNs (SNNs had less number of layers than DNNs, but each model in SNNs or DNNs had the same number of layers), Each model was trained independently on the naturalistic synthetic stereo images from the Monkaa dataset ^31^ using identical training protocols (see Methods for the details of the architectures and training scheme).

SNNs exhibited distinct learning dynamics across different binocular interactions, as reflected in their training and validation loss curves (Fig. 3b, the two upper-left panels). Although all SNN variants exhibited decreasing L1 loss during training, their validation performance remained poor (3-pixel accuracy ≤ 0.5; see Methods for the definition of 3-pixel accuracy; Fig. 3b, the two upper-right panels). Among the SNNs, the BEM exhibited the poorest validation performance (purple line), followed by the CMM (red line). While correlation-unconstrained architectures (Concat and Sum-diff) performed slightly higher validation accuracy and largely overlapped each other, yet their asymptotic validation performance remained close to 3-pixel accuracy of 0.5, meaning that nearly 50% of pixels in the predicted map deviates from the ground truth values by more than three disparity units. Correspondingly, the predicted disparity maps on the validation dataset generally appeared fragmented and noisy, indicating that shallow architectures, regardless of binocular interaction types, lacked representational capacity to extract reliable disparity information (Fig. 3c, upper panel).

In contrast, DNNs demonstrated substantially improved validation performance across all binocular interactions (Fig. 3b, lower panel). Despite their architectural differences imposed by the binocular interactions, they converged to similar L1 loss values and reached asymptotically around 0.85 accuracy. The Sum-diff architecture, however, converged at a slower rate. The predicted disparity maps were markedly more accurate than those generated by SNNs (Fig. 3c, lower panel).

In SNNs, correlation-constrained architectures (BEM and CMM) performed worse than correlationun-constrained networks (Concat and Sum-diff). This suggests that, under limited representational capacity, the imposed binocular interaction form might affect how features were represented across layers. Possibly, correlation-constrained SNNs formed a poor representational space that impaired the network’s performance. By increasing the representational capacity ^32^, however, DNNs incorporating correlation-based interactions achieved accuracy comparable to correlation-unconstrained models, or even slightly exceeding Sum-diff. This suggests that increased depth / representational capacity could compensate for the performance differences associated with the correlation mechanisms.

### Correlation-constrained models failed to capture key aspects of human depth perception in RDSs

Next, we conducted *in-silico* experiments to estimate each neural network model’s depth performance and its alignment with human behavior (Fig. 4a). Specifically, we fed each trained model with cRDSs, hmRDSs, and aRDSs across dot densities ranging from 10% to 90% in increments of 10%. The predicted disparity maps for cRDSs were used to train a linear SVM to classify crossed *vs.* uncrossed disparities. The trained classifier was then tested on disparity maps predicted for aRDSs and hmRDSs. This cross-conditioning decoding procedure resembled the multivariate analysis used in fMRI data. The SVM classification accuracy served as the model’s metric of depth performance. We then compared each network’s depth performance with human near/far discrimination across these RDSs.

Figure 4b shows representative disparity maps predicted by SNNs (upper panel) and DNNs (lower panel) for the three types of RDSs (target disparity = 10 pixels; dot density = 50%). For SNNs, the predicted disparity maps appeared noisy: relatively coherent for cRDSs, less clear for hmRDSs, and highly degraded for aRDSs. As a result, crossed and uncrossed disparity maps in aRDSs were not reliably distinguishable, regardless of binocular interaction type. In contrast, disparity maps generated by DNNs were markedly cleaner and more precise across RDSs. In particular, DNNs with Concat and Sum-diff interactions exhibited reversed disparity maps for aRDSs relative to cRDSs: uncrossed disparities (bluish) were predicted as near (reddish), while crossed disparities (reddish) were predicted as far (bluish).

Human behavioral performance is depicted in the upper rightmost panel of Figure 4c (highlighted in a yellowish rectangle). Participants exhibited near-perfect performance for cRDSs regardless of dot densities, gradually reduced accuracy for hmRDSs as dot density increased, and consistently below-chance performance, i.e., reversed depth, for aRDSs across dot densities. These behavioral patterns constituted the key aspects of human depth perception in RDSs, which served as the behavioral reference pattern for model comparison.

The left and middle panels of Figure 4c show model performance across RDS conditions. Correlation-based interactions in both SNNs and DNNs (outlined in blue) deviated substantially from the key behavioral signatures observed in humans for aRDSs and hmRDSs. In particular, these models exhibited relatively stable above-chance performance (0.5) for hmRDSs across dot densities and generally failed to produce systematic reversed-depth predictions for aRDSs, with the exception of the DNN implementing CMM at high dot densities.

Interestingly, the DNN with BEM interaction demonstrated the poorest alignment with humans (absolute mean difference, *AMD* = 0.2515 ± 0.000079; Fig. 4d; Fig. S2; Table S1), although they achieved validation accuracy comparable to other DNNs (Fig. 3b).

In contrast, the DNN incorporating the Concat binocular interaction showed a closer behavioral alignment with humans (AMD = 0.093; Fig. 4d; Fig. S2). It demonstrated decreasing performance with increasing dot density for hmRDSs and reversed-depth for aRDSs across a broad range of dot densities (30-90%). At low dot densities (10-20%), however, the model predicted above-chance performance for aRDSs, diverging from humans who consistently reported reversed depth across dot densities.

High validation performance on the Monkaa validation dataset (Fig. 3b) did not guarantee their behavioral alignment with human depth perception in RDSs (Fig. 4c). For example, the DNNs implementing CMM and Sum–diff binocular interactions achieved high validation accuracy (Fig. 3b), yet exhibited poor behavioral alignment with humans (Fig. 4c, d). Both networks predicted reversed-depth for aRDSs across dot densities (50 - 90% for CMM and 40 - 90% for Sum-diff). However, their performance for hmRDSs declined less steeply with increasing dot density compared to the DNN-Concat model. These discrepancies indicate that deep networks with different binocular interactions learned distinct disparity representations, and that certain aspects of human depth perception remain unresolved even in high performance deep hierarchical architectures.

Collectively, these results suggest two key points. First, among the tested architectures, deep hierarchical feedforward networks incorporating correlation-unconstrained computations could better align with human stereoscopic perception. Second, models constrained by explicit interocular correlation failed to capture the full behavioral patterns of human stereopsis. Additional constraints and/or learning objectives may therefore be required to fully account for the mechanisms underlying stereo processing in the human brain.

### Representational overlap differed across binocular-interaction architectures

The in silico psychophysics showed that correlation-constrained architectures, including BEM and CMM in both SNNs and DNNs, did not fully reproduce key signatures of human RDS behavior (Fig. 4c, blue outline; Fig. 4d). In contrast, the DNN-Concat architecture demonstrated the closest alignment with human behavior among the tested models (Fig. 4c, magenta outline; Fig. 4d). We therefore asked whether these behavioral differences were associated with differences in representational geometry. To do so, we analysed internal model representations and fMRI activity patterns using metrics inspired by superposition theory from AI interpretability research ^17^. Here, we used this framework operationally to quantify how separable or overlapping (representational overlap) the tested feature representations are across architectures and visual areas, rather than as direct evidence for the underlying mechanism.

We applied two complementary metrics to assess representational overlap. For SNNs, we computed feature dimensionality, which measures the effective dimensionality allocated to an individual feature within a layer’s embedding space ^17^ (see Methods, Eq. 2). The scores range from 0 to 1. Lower scores indicate that the feature shares more of its representational subspace with other features, suggesting greater representational overlap. Higher scores indicate that the feature representation is more separable from other features, indicating weaker overlap.

For DNNs and the human fMRI data, we quantified monosemanticity ^33^, which measures how selective an individual unit (or voxel) responds to a single stimulus feature or condition. Although monosemanticity and feature dimensionality are computed differently, they are conceptually related (see Methods). High monosemanticity suggests that a unit (voxel) is more specialized to process a single feature, suggesting weaker representational overlap among tested features. Lower monosemanticity suggests that the unit represents multiple features in the same embedding dimension, suggesting greater representational overlap among tested features. Crucially, monosemanticity can be computed both for DNN units and fMRI voxel responses, allowing a coarse comparison between biological and DNN representations, but not a direct equivalence between DNN units and fMRI voxels. This comparison cannot be evaluated using feature dimensionality analysis, which requires access to layer-to-layer weight matrices, which are unavailable in fMRI data (see Eq. 2 in Methods). We therefore used monosemanticity as an analogous, but indirect, measure of representational overlap in DNNs and human cortex. However, this comparison should be interpreted carefully. A 2-mm isotropic fMRI voxel pools signals from hundreds of thousands of neurons, so voxel-level monosemanticity should be interpreted as an aggregate measure of representational overlap in the voxel, not as evidence about the representational structure of individual neurons within that voxel. We simply used voxel-level monosemanticity for fMRI voxels and unit-level monosemanticity for DNN units to distinguish them.

The feature dimensionality analysis on SNNs differed across the binocular interaction types and layers (Fig. 5a). A two-way ANOVA revealed significant main effects of binocular interactions (*F* (3, 36) = 709.89*, p <* 0.001) and layer (*F* (3, 36) = 12127.56*, p <* 0.001), and a significant interaction between them (*F* (9, 108) = 245.53*, p <* 0.001). The feature dimensionality was highest in layer 1 and gradually declined as we ascended the layers. Pairwise comparisons between the binocular interaction types show that only CMM and BEM did not differ significantly from each other (the post-hoc Tukey tests, mean-difference = 0.0003, *p* = 1.0). However, both architectures exhibited significantly lower feature dimensionality across all layers than Concat and Sum-diff (BEM *vs.* Concat: mean-difference = 0.065, *p <* 0.001; BEM *vs.* Sum-diff: mean-difference = 0.102, *p <* 0.001; CMM *vs.* Concat: mean-difference = 0.066, *p <* 0.001; CMM *vs.* Sum-diff: mean-difference = 0.102, *p <* 0.001).

Lower feature dimensionality in correlation-constrained and limited representational capacity networks (SNNs) suggests that these architectures represented features with greater representational overlap than shallow architectures not constrained by explicit interocular correlation, such as Concat and Sum-diff. This difference was particularly pronounced in layer 3 where monocular left and right inputs first interact. Feature dimensionality did not differ significantly between BEM and CMM, given that they essentially come from the same computational family (both architectures represent features using a similar disparity-encoding strategy). The greater representational overlap in correlation-constrained SNNs could increase readout cross-talk and might contribute to poorer validation performance (Fig. 3b).

We next assessed monosemanticity as an indirect index of representational overlap in DNNs and the human visual cortex ^33^. When we grouped the data across RDS types, both humans and DNNs exhibited an increase in monosemanticity hierarchically (Fig. 5b). In humans, voxel-level monosemantic per feature increased from early to mid-higher brain areas (Fig. 5b, left panel). A repeated measures ANOVA revealed a significant effect of ROI on monosemanticity (*F* (7, 147) = 21.804*, p <* 0.001). A linear mixed model indicated a positive linear trend along V1-V2-V3-V3A-V3B-V7 (slope *β* = 0.317*, p <* 0.001), and V1-V2-V3-hV4-hMT+ (*β* = 0.457*, p <* 0.001) pathways. These suggest that disparity representations become more specialized (less entangled) at higher stages.

Likewise in the DNNs, unit-level monosemanticity increased with network depth, particularly in the Concat architecture (Fig. 5b, right panel). A repeated measures ANOVA revealed significant effects of binocular interaction (*F* (3, 36) = 17.975*, p <* 0.001), layer (*F* (18, 216) = 11.782*, p <* 0.001) and their interaction (*F* (54, 648) = 2.108*, p <* 0.001). A linear mixed model showed that the DNN-Concat model exhibited a significant positive trend relative to the BEM (*β_Concat_* − *β_BEM_* = 0.004*, p <* 0.001), while CMM and Sum-diff did not differ significantly from BEM (*β_CMM_* − *β_BEM_* = 0.0014*, p* = 0.148; *β_Sum_*_−_*_diff_* − *β_BEM_ <* 0.001*, p* = 0.69). Together, these observations suggest that both biological and artificial systems share a similarity in representing disparity: the disparity representation becomes more specialized along the processing hierarchy.

## Discussion

Binocular correlation models derived from single-cell responses have provided detailed accounts of the neural computations underlying biological stereopsis. These models successfully capture important aspects of local response properties of individual V1 neurons ^2,3,7–10,13,18,34^, especially the inverted disparity tuning for anticorrelated random-dot patterns ^11,12^. Many correlation-based accounts propose that stereo perception begins with correlation computations ^6,8–10,12^. If these local correlation-like signals directly formed the population codes used for perceptual inference, then reversed-depth perception (Fig. 2a), a psychophysical prediction of correlation models, should be reflected in V1 population response patterns. However, our fMRI multivariate analysis revealed that the perceptually aligned inverted representations were not linearly decodable in V1 but were detected in V3A (Fig. 2b, c). This dissociation raises a critical question: can correlation-based computations scale from local single-neuron responses to population codes that support human depth inference when binocular inputs are ambiguous?

### Multi-format stereo coding may better explain human depth perception

Our results suggest that local correlation-like signals may not directly form population representations that support human depth judgments under ambiguous inputs. One possibility is that aRDS-evoked disparity signals are present in V1 but organized in a representational geometry that is not linearly separable at the voxel scale. Responses to anti-correlated stimuli are not exactly the inverse of the correlated response profile; they are attenuated and not uniformly phase-shifted by 180^◦^. The tuning properties differ among cells, so that different neurons exhibit different response attenuation and phase shifts when responding to anticorrelated stimuli ^11,16^. Additionally, some V1 neurons with different RF profiles in the two eyes (phase disparity neurons ^15,34,35^) respond maximally to dissimilar features that do not correspond to a single physical feature in space (impossible stimuli ^18^). Such heterogeneous responses may preserve local disparity-related information, but after pooling within fMRI voxels, the representations of crossed and uncrossed disparities may overlap, even when cRDSs and aRDSs evoke comparable percent BOLD signal changes in V1 and other areas (Supplementary Fig. 3 in ref. ^22^). Thus, the undetected inverted representation from V1 voxel response patterns does not imply that V1 lacks anticorrelated disparity signals. Rather, those anticorrelated signals may be represented in a format that is not accessible to our linear decoder.

Complementing this fine-to-coarse-scale organizational account, our neural-network simulations provide a possible computational rationale for the discrepancy between single-neuron and population-level responses to anticorrelated patterns. Architectures explicitly constrained to correlation-like binocular interactions did not fully reproduce the key aspects of human depth judgments in RDSs, whereas architectures preserving monocular information before later integration aligned better with behavior (Fig. 4c, d). These results qualify other simulations demonstrating that pooling correlation-based neuronal activities in V1 can reproduce certain aspects of human performance ^7^. However, our results challenge the view that population representations exclusively encoded by correlation-based computations can support the full pattern of human depth perception.

Our representational analyses inspired by superposition theory in artificial networks ^17^ offer a helpful insight into why correlation-constrained networks did not fully reproduce the full pattern of human depth judgments. We quantified representational overlap to measure the degree of overlap among feature directions in activation space, which can increase the potential for cross-talk during readout. The analyses showed that architectures constrained by explicit correlation-like computations exhibited stronger representational overlap (Fig. 5a, b), which may reduce the accessibility of depth-relevant information to downstream readouts. This interpretation is consistent with the higher training losses observed in correlation-based shallow architectures (Fig. 3b; see ^17,36^). Simply scaling up our feedforward correlation-constrained architectures did not produce human-like depth perception (Fig. 4). One possibility is that units encoding many conjunctions of binocular features may form representations that are harder to learn and generalize when many features must be embedded in a limited-dimensional space ^37–39^. Another possibility is that early interocular multiplication discards monocular information that remains useful for later disparity inference. In the tested architectures, for example, anticorrelated monocular features can produce negative interocular products (*L* × *R <* 0) that are removed by rectification, potentially causing early information loss. This provides a possible explanation for why correlation-constrained networks did not facilitate generalization to novel stimuli (RDSs, particularly aRDSs and hmRDSs), and showed poor alignment with human behavioral data (Fig. 4c, d).

In contrast, models that preserved separate monocular features before later integration (particularly the DNN-Concat architecture) learned less entangled representations (Fig. 5a, b). This representational format was consistent with lower training losses in the shallow network (Fig. 3b) and a closer alignment with human behavior in the deep model (Fig. 4c, d). However, this analysis is diagnostic rather than causal: it shows an association between representational overlap and poorer model alignment with human behavior, but does not by itself establish that overlap alone causes the poor alignment.

Our findings, therefore, neither imply that V1 implements Concat-like processing, nor rule out correlation-based computations in early stereopsis. A more plausible account is that biological stereopsis may depend on a multi-format stereo coding strategy in which local correlation-like disparity signals are preserved alongside monocular and/or weakly integrated binocular representations ^40–42^ to be transformed or integrated later in the higher cortex. Encoding multiple binocular formats, rather than relying on correlation-like signals alone, may allow higher visual areas to retain the information that could be lost under early correlation-like transformations, thereby enabling the formation of population codes that better support perceptual depth. Such a hybrid account may explain why inverted population representations were detectable in V3A but not V1 (Fig. 2b, c), and why correlation-unconstrained deep networks more closely matched human depth judgments (Fig. 4c, d).

### Voxel-level monosemanticity progressively increases along the visual hierarchy

The voxel-level monosemanticity analysis, inspired by monosemanticity metrics in artificial networks ^33^, showed increasing disparity-specific organization along the human visual hierarchy (Fig. 5b). Lower voxel-level monosemanticity in early visual cortex suggests that feature representations in early stages are consistent with greater representational overlap at the voxel level than those in higher cortical areas. This result is expected because V1 neurons are selective to a diverse set of low-level features, such as disparity, contrast polarity, orientation, complex pattern, and combinations of them ^43^. Many of these raw sensory signals might intermix within an fMRI voxel, obscuring disparity-relevant representations even if they are present at finer neural scales.

In later visual areas, higher voxel-level monosemanticity suggests a reduced representational overlap among disparity-relevant features. In V3A, this organization was also aligned with the perceptual signature of reversed depth, suggesting that disparity representations might be transformed from a local sensory format into re presentations that were more spatially organized and behaviorally relevant. This reasoning is compatible with the view that early sensory representations reflect a relatively faithful copy of sensory inputs whereas brain representations in later areas are transformed into behaviorally relevant abstractions ^44^.

Applying the same monosemanticity analysis to DNN-Concat revealed a qualitatively similar trend. Disparity specificity increased across layers, in parallel with the increase observed across human visual areas (Fig. 5b). In contrast, architectures constrained by explicit correlation-like computations did not show a significant positive trend across layers. This suggests that architectures that exclusively encode disparity using correlation-like computations can learn the training objective, yet fail to learn layer-wise representational spaces that align with human brain representations and behavior, even when their validation performance approaches that of correlation-unconstrained deep networks (Fig. 3b).

However, a cautionary point needs to be added to this voxel-level monosemanticity interpretation. The metric used for single fMRI voxels differs from that in single DNN units due to differences in spatial resolution. A single voxel contains hundreds of thousands of neurons and can be considered as a weighted linear combination of their activity shaped by spatiotemporal filtering properties ^45^. Consequently, the high voxel-level monosemanticity observed in V3A may not straightforwardly reflect the monosemanticity of individual units. monosemanticity in the higher cortex should be interpreted as a macro-scale topographic clustering at the population level ^46^.

### Anticorrelated responses in hMT+

Recent studies using population receptive field (pRF) methods have shown population disparity tuning profiles for anticorrelated dot patterns across human visual cortex ^47,48^. In particular, Parker et al., ^48^ reported that hV5/MT shows a signature of the binocular energy model predictions, i.e., the inversion of population disparity profile between correlated and anticorrelated stimuli (Fig. 4 in ref. ^48^). However, the population tuning curves for aRDSs in that study were obtained by averaging vertex-wise disparity-pRF estimates and then normalizing the resulting curves. Therefore, the responses to anticorrelated patterns should not be interpreted as equal to that of correlated stimuli. Indeed, anticorrelated responses were generally weaker (the amplitude ratio between aRDSs and cRDSs was reported 0.258 for human pRF ^48^) and less structured than correlated responses (see Fig. S2.1 in ^48^ and Fig. 6b in ^47^).

Nevertheless, we do not interpret the findings of Parker et al., ^48^ as contradictory because our data also shows that hMT+ contains voxels with a strong disparity specificity to aRDSs (Fig. S3). Thus, the multivariate analysis results in Fig. 2b and c do not claim the lack of responses to anticorrelated stimuli in hMT+. Rather, they indicate that hMT+ did not show reliable population representations consistent with reversed depth perception for anticorrelated dot patterns.

A likely source of this difference is methodological. Parker et al., ^48^ used a different unit of analysis. The authors used BOLD responses sampled from surface-vertices selected based on modified pRF methods, whereas we used voxels selected according to a standard GLM analysis. Specifically, Parker et al., ^48^ employed pRF pipeline extended to depth to obtain surface vertex data whose BOLD time course signals were sufficiently explained by the disparity-pRF model and whose fitted parameters were physiologically plausible. This procedure is optimized for identifying cortical responses that are specifically sampled from the fMRI data whose pRFs fall within the stimulating region in the visual field.

In contrast, our multivariate analysis selected the top 250 task-sensitive voxels according to the GLM analysis comparing all stimulus conditions against fixation (see Methods). This voxel selection criteria was used to provide feature dimensions for multivariate decoding, but it was not optimized for isolating voxels whose spatial pRFs fell within the stimulating regions (disparity-defined apertures). Although we could not tailor stimulus position for every voxel, we selected the top task-sensitive voxels which were engaged in our stimuli. Nevertheless, it would be interesting to use voxels selected based on the criteria used in Parker et al., ^48^ for multivariate analysis to see if the information decoding would differ from ours.

### Study limitations

We did not incorporate temporal dynamics of stereo processing into our models, which have been shown to influence depth perception, particularly for anticorrelated stimuli ^21^. To assess the potential role of temporal dynamics behaviorally, we conducted additional human psychophysics comparing depth-discrimination performance for static and dynamic RDSs (Fig. S1). Participants could perceive reversed depth for dynamic aRDSs, but not for the static case, suggesting that temporal signal integration may contribute to the perceptual interpretation of anticorrelated disparity signals in humans.

The effect of absence of temporal signal integration mechanisms in our models is reflected in their responses to aRDSs. Although DNN-Concat showed less entangled disparity-related representations than correlation-constrained models, its poor alignment at sparse dot densities may indicate the limit of DNN as static frame models (Fig. S2). Incorporating temporal dynamics may therefore be necessary for future models to more fully account for human stereo perception under static and dynamic viewing conditions.

## Conclusion

We investigated whether correlation-based disparity computations, which explain important local response properties of binocular neurons, scale into population-level representations that support human depth inference under ambiguous binocular input. Our results show that brain representations consistent with reversed depth perception were explicit in V3A but not detected at voxel scale in V1. Neural networks constrained to explicit correlation-like interactions failed to capture the full pattern of human depth judgments. These suggest that population codes with exclusive correlation-like computations may not directly support human-like stereopsis.

Our study provides a new insight into why population codes encoded exclusively by correlation-like computations may not fully support human depth perception. In the tested models, architectures relying solely on correlation-like interactions exhibited stronger representational overlap and demonstrated poor alignment with human behavior, whereas architectures preserving monocular information before later integration better matched human depth judgments. These simulations do not rule out correlation-based computations in early stereopsis. Rather, they suggest that biological stereopsis may require additional representational formats beyond direct pooling of local correlation signals. We therefore propose a multi-format stereo-coding account in which correlation-like, monocular, and weakly integrated binocular information are preserved and transformed along the visual hierarchy to support perceptual depth inference under ambiguity.

## Methods

The fMRI dataset and accompanying psychophysical measurements shown in Figure 2 were previously acquired and reported ^22^. In the present study, we re-analysed these data using new and independent analyses to address a distinct research question about whether population representations encoded by correlation-based computations can support human depth perception. Specifically, the multivariate cross-decoding and searchlight analyses reported in Fig. 2b, c were newly performed for this study and were not included in our previous paper ^22^. We further performed new neural network modeling in which binocular-interaction layers were incorporated into shallow and deep architectures, and collected a new psychophysical dataset for the model-comparison analyses shown in Fig. 4. Finally, we applied a representational analysis inspired by superposition theory from AI-interpretability research to characterize the internal model representations shown in Fig. 5.

### Participants

Twenty-five participants (five females, 20–40 years, mean = 24.5, SD = 4.37) took part in the combined fMRI and psychophysics experiments (Fig. 2). Data from three male participants were excluded: two did not complete the psychophysics experiment, and one exhibited excessive head motion during fMRI scanning. An additional 33 participants (13 females, 18–44 years, mean = 23.67, SD = 4.66) were recruited for a second psychophysics experiment (Fig. 4). Finally, 21 participants (10 females, 20–60 years, mean = 26.26, SD = 10.12) participated in the third psychophysics experiment (Fig. S1). All participants had normal or corrected-to-normal vision. All ethical regulations relevant to human research participants were followed. All procedures were performed in accordance with the ethical standards stated in the Declaration of Helsinki 2008 and approved by the Ethics and Safety Committee at the Center for Information and Neural Networks (CiNet), National Institute of Information and Communications Technology (NICT). Written and oral informed consent was obtained from all participants prior to participation.

### Task and visual stimuli

For the fMRI experiments, visual stimuli were generated in MATLAB (MathWorks Inc.) with psychtoolbox ^49^. Stimuli were back-projected onto a custom translucent screen (19.7*°* × 15.8*°* visual angle, Kiyohara Optics, Inc., Tokyo, Japan) positioned at the rear of the MRI bore. Stereo presentation was achieved with two LCD projectors (1920 × 1200 pixels, 60 Hz refresh rate; WUX4000, Canon, Tokyo, Japan), each fitted with a distinct polarizing filter to separate the left and right images. Projector luminance values were gamma-corrected and matched using a MATLAB toolbox publicly available at https://github.com/hiroshiban/Mcalibrator2^50^ and a colorimeter (Konica-Minolta, CS-100A). The filtered left and right RDS images were carefully aligned by superimposing geometric test patterns (rectangles, circles, and grids of different colors) using a custom calibration tool (https://github.com/hiroshiban/BinocularDisplayChecker) prior to the experiment. Participants viewed the stimuli through a tilted front-surface mirror at a distance of 96 cm while wearing polarized glasses (KGA04-B, Kiyohara Optics, Inc., Tokyo, Japan). They were instructed to maintain fixation on a central white square (0.6*°* visual angle) and to perform a detection task to maintain their attention ^51^. This task involved identifying the position of a randomly appearing white vertical bar briefly presented within the square for 250 ms. Participants pressed a left or right button depending on the bar’s position relative to the upper nonius line, or refrained from responding if no bar appeared.

During the detection task, participants viewed dynamic RDSs (dot density = 25%, dot refresh rate = 30 Hz) painted on a mid-gray background ^52^. Stimulus timing is described in the *fMRI stereo experiments* section. The RDSs contained equal proportions of black and white dots (50% each; diameter = 0.14*°*). Luminance was set to 128 *cd/m*^2^ for white dots, 64 *cd/m*^2^ for the mid-gray background, and 0.39 *cd/m*^2^ for black dots. Thus, the average of the dot luminance was closely matched to the gray background, minimizing luminance contrast change during stimulus onsets and offsets.

The stimuli comprised a circular background RDS (14*°* diameter) with four identical smaller circular RDSs (4*°* diameter each) presented in the four quadrants, centered at 2.5*°* horizontally and vertically from the center of the background RDS (Fig. 1a). These four target RDSs were presented with crossed (–0.2*°*) or uncrossed (+0.2*°*) disparities. The background RDS was placed on the fixation plane (0*°* disparity) and was always binocularly correlated (i.e., black and white dots on the left image were consistently paired with black and white dots on the right image, respectively). The four target RDSs varied in dot correlation: correlated RDSs (cRDSs, correlation = +1), half-matched RDSs (hmRDSs, correlation = 0), and anticorrelated RDSs (aRDSs, correlation = –1). cRDSs comprised dots of identical contrast in the two eyes; aRDSs had opposing contrast dots—black dots in the left image paired with white dots in the right image, and vice versa—; and hmRDSs comprised an equal mix of anticorrelated and correlated dots, resulting in a zero net correlation. The dot density was 25%, and the dot refresh rate was 30 Hz.

### fMRI image acquisition

Functional BOLD responses were acquired on a Siemens MAGNETON Trio 3T Scanner at NICT, CiNet imaging facility (Suita, Osaka, Japan) using a 32-channel phased-array head coil. Since we were primarily interested in the occipito-parietal and occipito-temporal cortex and participants wore polarized glasses during the scanning, only the occipital part of the head coil was used. Functional images were collected with the following parameters: voxel size = 2 mm isotropic, repetition time (TR) = 2 s, 78 slices, multiband factor = 3, field of view (FOV) = 192 × 192 *mm*^2^, and flip angle = 75*°*. Each functional run comprised 208 volumes. The Multiband EPI sequence was provided by the University of Minnesota (under a C2P agreement, https://www.cmrr.umn.edu/multiband/. High-resolution T1-weighted anatomical images covering the whole brain were acquired for each participant (voxel size = 1 mm, 208 slices, TR = 1900 ms, echo time = 2.48 ms, FOV = 256 × 256 *mm*^2^, flip angle = 9*°*). These anatomical scans were used for cortical surface reconstruction and precise coregistration of functional and structural images.

### Retinotopic mapping and cortical preprocessing

We performed a separate session for localizing regions of interest (ROIs). ROIs (V1, V2, V3, V3A, V3B, V7, hV4, and hMT+) were defined using standard retinotopic mapping procedures ^53–56^. Stimuli consisted of clockwise (CW)/counterclockwise (CCW) rotating wedges, as well as expanding and contracting concentric rings. Ten participants completed a single run of CW rotating wedge stimuli on the same day as the main fMRI stereo experiments. The remaining 12 participants performed seven runs on a separate day: two runs with CW rotating wedges, two with CCW rotating wedges, one run with expanding rings, one with contracting rings, and one with dot patterns alternating between motion and stationary for localizing hMT+ ^57^. Stimulus codes are available at https://github.com/hiroshiban/Retinotopy. Cortical reconstruction was performed in BrainVoyager QX ver 2.8 (BrainInnovation B.V, Maastricht, The Netherlands) following established procedures ^58^. Briefly, noncortical structures (skull) were removed, anatomical scans were transformed into Talairach space, flattened surface representations of the two cerebral hemispheres were generated.

### fMRI stereo experiments

We employed a block design in which each functional run comprised 24 blocks of 192 trials, began and ended with a 16 s stimulus-off period (mid-gray background with a fixation marker). Each block of 8 trials lasted 16 s and consisted of alternating periods of 1 s stimulus-on and 1 s stimulus-off (mid-gray background). To minimize potential brain responses to luminance contrast changes between the stimulus and interstimulus periods, we carefully matched the mean monocular luminance of all RDS types and blank screen to a mid-gray background.

Each functional run lasted 416 s in total (16 s stimulus-off + 24 blocks × 8 trials-per-block × 2 s-per-trial + 16 s stimulus-off). Within a block, only one of the six stimulus conditions (cRDS, hmRDS, and aRDS, each with either crossed or uncrossed disparity) was presented. Stimulus order was randomized across blocks to prevent sequence effects. A 10 s “dummy” scan was inserted at the beginning of each run to avoid transient signal drifts.

In the main experiment, participants completed different numbers of runs. Four participants accomplished 11 runs, seven completed 10 runs, three completed nine runs, five completed eight runs, two completed six runs, and one completed four runs.

### Psychophysics experiments

We conducted three psychophysics experiments. All psychophysics experiments employed a two-alternative forced choice (2AFC) task in which participants judged near / far from the RDS stimuli. Prior to the main experiments, their stereo-acuity was measured and only participants with good stereo-acuity were included. Each trial involved two sequential presentations of the same RDS type (aRDS, hmRDS, or cRDS): one with crossed disparity (–0.2*°*) and the other with uncrossed disparity (+0.2*°*), presented in random order. Each stimulus was presented for 1 s, separated by a 1 s mid-gray background interval, yielding a trial duration of 3 s. After the stimulus presentation, participants indicated whether the first or second stimulus appeared closer by clicking the left or right mouse button, respectively. They then pressed the space bar to proceed to the next trial, with no time limit for their response. To minimize the adaptation effect across trials, random disparity jitter up to ±1 arcmin was added to each dot.

We expected two possible perceptual outcomes: “forward-depth” and “reversed-depth”. In the forward-depth scenario, uncrossed-disparity would be perceived as far, and crossed-disparity as near. In the reversed-depth scenario, depth would be perceived in the opposite direction to forward-depth, i.e., uncrossed-disparity would be perceived as near, and crossed-disparity as far (see Fig. 1b). For participants who experienced reversed-depth in aRDSs, the expected proportion correct of the response would be below chance.

The first experiment was aimed to measure depth performance in dynamic aRDSs, hmRDSs, and cRDSs at a fixed dot density of 25% and dot refresh rate at 30Hz (Fig. 2). The stimuli were identical to those used in the fMRI experiment but presented on an LCD monitor (60 cm × 44.5 cm, 3840 × 2160 pixels, 120 Hz refresh rate). Participants viewed the stimuli through NVIDIA 3D Vision LCD shutter glasses at a viewing distance of 57 cm. Participants’ responses were recorded via computer mouse. Each stimulus was repeated eight times per run. Most participants accomplished four runs, only one participant completed three runs.

The second experiment assessed depth performance for dynamic aRDSs, hmRDSs, and cRDSs as a function of dot density (Fig. 4c). The dot refresh rate was kept constant at 30 Hz while systematically varying dot density from 10% to 90% in increments of 10%. Each stimulus was repeated five times per run. All participants completed five runs.

The third experiment aimed to compare the depth performance under static (dot refresh rate = 0 Hz) and dynamic conditions (dot refresh rate = 30 Hz) at a fixed dot density of 25% (Fig. S1). Each stimulus was repeated five times per run. Ten completed six runs, six completed four runs, five participants completed three runs.

### Cross-decoding analysis

We performed cross-decoding (multivariate) analysis on our fMRI data to identify neural correlates of reversed depth perception in the human brain. Specifically, a linear classifier (support vector machine, SVM) was trained on brain activity patterns evoked by crossed and uncrossed disparities in cRDSs and tested on patterns evoked by aRDSs, and *vice-versa*. The rationale was based on the inverse relationship between disparity tuning for cRDSs and aRDSs, where a population of disparity-selective neurons invert their disparity tuning profile when presented with opposite dot polarity (binocular correlation is inverted) ^11,22,26^. Accordingly, classification performance significantly below chance would indicate the neural signatures associated with reversed depth.

Voxels for this analysis were selected by applying a general linear model (GLM) that assesses the contribution of each experimental condition to voxel time course variance. We focused on voxels where the effect size contributed by all stimulus conditions was more significant than the baseline (fixation) condition. The GLM analysis for the whole-brain was done for each participant. The design matrix included regressors for the experimental conditions (3 RDS types × 2 disparity signs), six motion parameters (head movements: translations in XYZ axes and rotations in pitch, roll, and yaw), and a constant term as regressors. The regression coefficients (GLM beta values) were derived from the least-squares solution to the linear equation. We then computed the *t*-statistic for each voxel within the predefined brain regions with a contrast vector of all stimulus conditions *vs.* fixation. The top 250 task-sensitive voxels for each individual were selected in descending order of *t*-values, and served as the feature dimensions for cross-decoding.

Cross-decoding analysis was implemented in Python using the scikit-learn library (https://scikit-learn.org/stable/), with a linear kernel. The regularization parameter C was set to 1, and default settings were used for other parameters. For each participant, we cross-validated the fMRI dataset for each ROI with a leave-one-run-out scheme to evaluate generalization. Cross-decoding accuracies for each ROI were then averaged across participants.

### Searchlight analysis

We performed a group-level searchlight analysis ^27^ to confirm that our predefined ROIs, obtained from independent retinotopic scans, encompassed the essential brain regions involved in reversed depth perception. For each participant, we applied a linear SVM using a leave-one-run-out cross-validation scheme to cross-decode local patterns of brain activity (cRDS *vs.* aRDS, *vice versa*) within a sphere (radius = 6 mm), traversing across the entire cortex in Talairach space.

This procedure generated information maps highlighting regions where local brain activity patterns evoked by crossed and uncrossed disparities in anticorrelated stimuli were inversely related to those of correlated stimuli, and *vice versa*. The *t*-statistics for the cross-decoding accuracies were computed against chance level performance (0.5). Significant clusters of voxels were identified by thresholding them at *p <* 0.005 (one tailed, uncorrected). The resulting searchlight map was overlaid onto a representative individual’s brain. Both searchlight and ROI-based analyses were conducted independently and complemented each other without circularity (”double-dipping”) in voxel selection. Accordingly, the searchlight results (Fig. 2c) were consistent with the ROI-based results (Fig. 2b) and do not alter our main findings or conclusions.

### Simulation of disparity processing with shallow and deep neural networks

We implemented shallow and deep convolutional neural networks to predict disparity maps from stereo image pairs. The shallow convolutional neural network was inspired by the Goncalves and Welchman model ^19^, but differed from the original implementation in two important ways. First, unlike the original model, which classified disparity signs, our shallow networks were trained to predict a continuous binocular disparity value at each pixel. Second, we did not initialize the networks with Gabor filters or impose any prior assumptions about receptive-field structure. This allows all convolutional filters to be learned directly from the training data.

To make the shallow model comparable to modern stereo architectures, we incorporated the cost volume tensor, following the design from the Geometrical and Contextual network (GC-Net ^59^). Concretely, our shallow binocular models can be regarded as the scaled-down version of GC-Net, allowing the models to learn disparity statistics from stereo inputs with a limited representational depth. Details of the architecture are shown in Fig. 3a and described below in *Shallow binocular neural network (SNN) architecture*.

For DNN, we implemented GC-Net ^59^ that has been shown to share a similar disparity representational transformation with the human brain ^22^. Architectural details are provided in Fig. 3a and described below in *Deep binocular neural network (DNN) architecture* for the details.

#### Shallow binocular neural network (SNN) architecture

Each SNN comprised three main modules: a feature extractor, a disparity cost volume, and a disparity regressor (Fig. 3a). The feature extractor module contained two monocular streams (one for each eye) with shared weights. Each monocular stream consisted of two 2D convolutional layers, each followed by a batch normalization layer and a ReLU nonlinearity. The cost volume was a 4D tensor that represented the concatenation of feature maps from one eye with spatially shifted maps from the other eye. The disparity regressor module comprised a 3D convolutional layer, a 3D deconvolutional layer, and a soft argmin operator (Eq. 1) to produce continuous disparity map.

The network received RGB stereo image pairs as input (three channels per image). In each monocular stream, the first convolutional layer applied 32 filters with kernel size 3 and stride 2, mapping the input from 3 to 32 channels. The second convolutional layer applied 32 filters with kernel size 3 and stride 1, preserving the channel dimensionality. Thus, each monocular feature extractor produced a feature map with 32 feature channels.

The left and right monocular feature maps from the feature extractor module were then concatenated along the channel dimension according to the specified binocular interactions (see *Binocular interactions*) before constructing the cost volume tensor. Specifically, for each disparity level, the feature map from one eye was concatenated with the corresponding spatially shifted feature map from the other eye, following the standard cost-volume formulation ^59^. This procedure was repeated across all candidate disparities, yielding a four-dimensional cost-volume tensor with dimensions [2*C, N_d_, H, W*], where *C* = 32 is the number of feature channels, *N_d_* = 192 is the number of candidate disparities, and the disparity set ranged from −96 to 95 pixels in 1-pixel increments.

The cost volume was processed by a 3D convolutional layer to learn feature representations from the height, width, and disparity dimensions. The 3D convolutional layer had 32 filters with a kernel size 3 and stride 1, mapping the input channels from 64 to 32. We then applied a single 3D transposed convolution (deconvolution), with kernel size 3 and stride 2, reducing the channel dimension from 32 to 1.

Finally, the output of the 3D transposed convolution was converted into a continuous disparity map using a soft argmin operation defined in Eq. 1. Specifically, a softmax was applied along the disparity dimension to obtain a probability distribution over candidate disparities, and the expected disparity for each pixel was computed as:

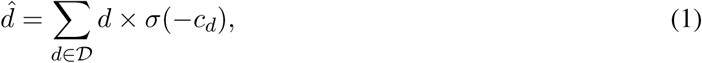

where D = {−96, −95*, …,* 95} denotes the candidate-disparity set, *c_d_* represents the predicted cost at disparity *d*, and *σ*(·) denotes the softmax operation.

#### Deep binocular neural network (DNN) architecture

We used GC-Net ^59^ as our DNN, which consisted of three main modules: a feature extractor, a disparity cost volume tensor, and a disparity regressor. The detail of the GC-Net architecture is provided in its original paper ^59^ and Fig. 3a for its schematic.

The feature extractor module contained two monocular streams, one for each eye, with shared weights. Each stream consisted of 18 2D-convolutional layers, each layer was followed by a batch normalization layer and a ReLU nonlinearity (except for the 18*^th^* layer). The disparity regressor consisted of 14 3D-convolutional layers, 5 3D-deconvolutional layers, and a soft argmin operator. Layer dimensionalities followed the original GC-Net implementation ^59^.

Cost-volume construction and disparity regression followed the same principles as in the SNNs, except that the DNN contained substantially more 2D and 3D processing stages. This allowed us to test whether representational depth altered the effect of different binocular-interaction assumptions.

#### Binocular interactions

To assess how different binocular-computational assumptions affect neural networks in learning the statistics of disparity from naturalistic stereo inputs, we tested four types of binocular interactions: two correlation-constrained and two correlation-unconstrained interactions (Fig. 3a). We incorporated correlation-constrained interactions to explicitly introduce correlation-like computations into the network (Fig. 3a). These included the Binocular Energy Model (BEM ^2^) and the Cross-correlation and Matching Model (CMM ^5,6,22^).

The BEM-inspired network combined the monocular energy and the interocular terms [(*L*^2^ + *R*^2^), (L × R)]. Here, L and R denote the convolutional layer outputs from the left and right convolutional streams, respectively. The CMM-inspired combined the interocular terms with their rectified counterparts [(L × R), ReLU (L × R)].

As for correlation-unconstrained networks, we implemented two binocular interactions that do not explicitly compute interocular correlation. These include Concat (concatenation) and Sum-diff (summing-differencing channels ^29,30^). The Concat represented a simple feature concatenation that combined the left and right monocular features as [L, R]. The Sum-diff combined the summing and differencing terms [L + R, L - R]. These four interaction rules were inserted at the same stage of the network architecture, allowing us to compare architectures with explicit correlation-like constraints against architectures in which binocular feature combinations were learned without such constraints.

#### Training procedure

All networks were trained on the synthetic Scene Flow Monkaa dataset, which provides precise ground truth disparity maps ^31^. To avoid train–validation leakage, we first divided the original stereo image pairs into separate training and validation sets. Random crops were then sampled only within each split, so that no crop from a validation image came from an image that was also used for training. The original stereo images (540×960 pixels) were cropped to patches of size 256×512 pixels, with the cropping locations were uniformly randomized across the entire images. Crop locations were sampled uniformly across the valid image region, allowing the networks to observe different spatial regions of each image across epochs.

To allow the networks to represent both positive and negative disparities, the disparity values in the Monkaa dataset were shifted by a constant horizontal offset. The offset was set to twice the mean ground-truth disparity of the Monkaa training set (2 × 44 = 88 pixels). The same disparity transformation was applied consistently to the stereo images and their corresponding ground-truth disparity maps. A batch size of four was used for both training and validation.

All models were implemented in PyTorch (version *>* 2.8) and trained for 10 epochs. Each architecture was trained independently with 13 random seeds: 1618, 11364, 16476, 27829, 35154, 35744, 36675, 43798, 55826, 59035, 65190, 82220, and 94750. Optimization was performed using the AdamW optimizer with a cosine learning-rate schedule (warm-up steps = 250, min learning rate = 6 × 10^−5^, max learning rate = 6 × 10^−4^). We applied gradient clipping with a maximum norm 0.1 to stabilize training. The training was accelerated on a GPU NVIDIA RTX 5090 (NVIDIA, California, USA), with compile mode “reduce-overhead”.

To decrease the likelihood of the network memorizing input examples, we applied data augmentation during training only. Data augmentation was not applied during validation, resulting in lower validation loss (higher accuracy) than training loss in the learning curves. The augmentation pipeline included random vertical shifts and rotations (`RandomShiftRotate`, `always apply=True`, other parameters at their default values), random RGB-channel intensity shifts (`RGBShiftStereo`, `p asym=0.3`, other parameters at their default values), additive Gaussian noise (`GaussNoiseStereo`, `always apply=True`, `p asym=0.3`, other parameters at their default values), and random brightness and contrast changes (`RandomBrightnessContrastStereo`, `always apply=True`, `p asym=0.5`, other parameters at their default values). In addition, left and right images were swapped with probability 0.5. When the left image served as the reference view, the corresponding left-view disparity map was used as the training target (ground truth). When the right image served as the reference view, the corresponding right-view disparity map was used as the ground truth.

Model performance was quantified using the L1 loss (SmoothL1Loss in PyTorch), the mean absolute error between the predicted disparity map and the ground truth. Training loss was recorded every 100 iterations and averaged over 200 iterations for visualization of the learning curves. We selected the model’s parameters based on the lowest validation loss.

As a complementary performance metric, we calculated 3-pixel accuracy. This was defined as the proportion of valid pixels for which the absolute difference between the predicted disparity and the ground-truth disparity was less than or equal to three disparity units.

### Measuring depth judgment performance for SNNs and DNNs

We conducted an in-silico psychophysics experiment to quantify depth judgment performance in SNNs and DNNs. RDSs were generated with variation in dot correlation (*T* = {aRDS, hmRDS, cRDS}, where *T* is the set of RDS types) and dot densities (*S* = {0.1, 0.2*, …,* 0.9}, where *S* is the set of dot densities). We generated 1000 RDS images for each combination of dot density and dot match level. These RDSs were presented as binocular inputs to our shallow and deep models.

For each dot density, we trained a linear SVM on the predicted cRDS disparity maps to classify near or far depth planes. Then, we tested the trained classifier on the predicted disparity maps from aRDSs and hmRDSs. The classification accuracy serves as the measure of the depth performance for aRDSs and hmRDSs at each dot density.

### Absolute mean difference (AMD) between human and AI psychophysics

To evaluate the degree of alignment between human and neural networks, we calculated the absolute mean difference (AMD) between their psychophysical performance. Concretely, we calculated the absolute of mean difference between human and model depth performance for each dot density and dot match level. We performed bootstrap resampling with replacement (1000 bootstrap resampling and 1000 iterations) and averaged the AMD across bootstrap samples (Fig. S2 and Table S1). Then, we averaged the bootstrapped AMD values across dot density and dot match level to obtain the alignment score, which was reported in Fig. 4d.

### Feature dimensionality

To understand why models behavior differed across binocular-interaction architectures, we analyzed the geometry of their internal representations using a diagnostic measure inspired by superposition theory from AI-interpretability research ^17^. In this framework, features are treated as directions in activation space. Specifically, these feature directions were those in experimentally defined stimulus conditions used in our analyses, rather than to a latent semantic feature learned independently by the model. We used this framework as an operational tool for quantifying the degree of representational overlap, not as evidence that biological circuits or artificial networks necessarily implement the same coding scheme. We used this metric to measure the degree of representational overlap in SNNs.

The superposition theory posits that a neural network represents features in superposition when it stores far more features than the available dimensions (units) in their activation space ^17^. If features were strictly orthogonal, a model with *m* neurons could only represent *m* unique features. However, when the model needs to encode *N* features in *m*-dimensional space with *N > m*, it cannot assign each feature its own orthogonal direction. Instead, features must share representational space, allowing feature directions to overlap “almost-orthogonally”. This sharing introduces interference (representational overlap) where multiple features compete for the same representational directions, so activating one would inevitably perturb others. In artificial networks, this overlap is related to polysemanticity, in which individual units contribute to multiple features ^17^. A related, but not identical, concept in neuroscience is mixed selectivity, in which neurons respond to combinations of multiple features ^60–62^, supporting flexible linear decoding for complex behavioral tasks ^39^. Mixed-selectivity neurons have been observed across brain areas: V1^43,63–65^, V4^65^, prefrontal cortex (PFC) ^60^, posterior parietal cortex (PPC) ^66,67^.

A metric used to quantify the degree of representational overlap is feature dimensionality. Feature dimensionality measures the effective fraction of embedding (representational) dimensions allocated to a given individual feature ^17^. It can also be understood as feature capacity: the fraction of a dimension allocated to a feature *i* ^68^. The values are bounded within a range of 0 (not learned) and 1 (dedicating a dimension to a feature). A value close to 1 indicates that the feature is nearly orthogonal to other features, occupying an almost dedicated dimension in the representational space. A smaller value signifies that many features share similar directions, reflecting a stronger superposition (representational overlap) which gives rise to polysemantic neurons. When features are not learned at all, then the feature dimensionality becomes 0.

Concretely, feature dimensionality of the *i*-th feature *D_i_* is defined as follows ^17^:

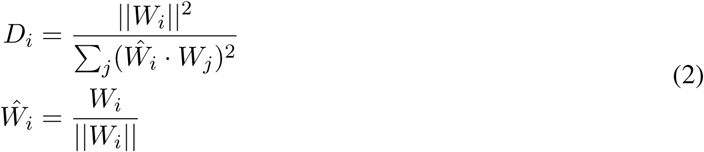

where *W_i_* ∈ R*^m^* represents the weight vector of the *i*-th feature in an *m*-dimensional embedding space; *Ŵ_i_* denotes its normalized vector. The squared norm of feature vector ||*W_i_*||^2^ in the numerator indicates how strong feature *i* is represented (representation strength). The denominator is called the *superposition index*, indicating how many features share their dimension with feature *Ŵ_i_* that is embedded in the space by projecting all feature vectors onto the direction of *Ŵ_i_*.

### Monosemantic score

For both fMRI data and DNN models, we quantified the degree of representational overlap using a monosemantic score adapted from ^33^. This score measures the extent to which an fMRI voxel or a DNN unit responds predominantly to a single tested feature, rather than responding broadly across many tested features. As in the feature-dimensionality analysis, “feature” here also refers to an experimentally defined stimulus condition used in our analyses, rather than to a latent semantic feature learned independently by the model. We therefore used the term monosemantic score operationally not in the full mechanistic sense used in AI-interpretability research ^33^. Because it is impossible to know the whole latent features represented by the brain, our analysis was restricted to stimulus variables that were physically defined, experimentally controlled, and matched across model and fMRI datasets. We simply refer to the monosemantic score as voxel-level monosemanticity for fMRI voxels and unit-level monosemanticity for DNN units.

The monosemantic score as a measure of representational overlap may share a similar interpretation with feature dimensionality. While feature dimensionality measures the degree of representational overlap in activation space, the monosemantic score measures the overlap at the level of individual response units. The monosemantic score is bounded between 0 and 1. A high score indicates that most of a voxel’s or unit’s is dedicated to a single stimulus-defined feature. A low score indicates that the response is distributed across multiple tested features. Thus, lower monosemantic scores are consistent with greater representational overlap at the voxel or unit level, although they do not directly measure geometric overlap among feature vectors.

Let *T* = {aRDS, hmRDS, cRDS} be the set of RDS types, *S* = {0.1, 0.2*, …,* 0.9} be the set of dot densities, and *D* = {near, far} be the set of disparity signs. We define the feature set 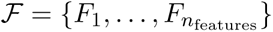 whose elements are ordered triples (*t, s, d*) in *T* × *S* × *D* = {(*t, s, d*) : *t* ∈ *T, s* ∈ *S, d* ∈ *D*}. Thus, the number of features *n_features_* = |F | = |*T* | × |*S*| × |*D*| = 3 × 9 × 2 = 54, where |*X*| denotes the cardinality of *X*.

For unit or voxel *i*, let *h_i_*(*F_j_*) be its activation to feature *F_j_*. For convolutional layers, we treated each output channel as a unit by averaging the feature map over spatial locations. Thus, for 2D convolutional layers, the number of units for a layer is given by *N* = *n*_feature−channels_, while for 3D convolutional layers, *N* = *n*_feature−channels_ × *n*_disparity−channels_.

The monosemantic strength measures how monosemantic a unit is: the degree to which a unit’s activation is specific to a single stimulus feature or condition ^33^. Specifically, the monosemantic strength is expressed as follow:

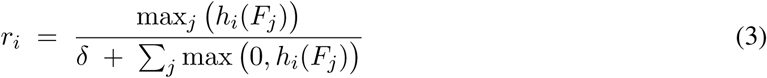

where *r_i_* corresponds to the strength of the activation of unit *i* to its most strongly activating feature *F_j_*divided by the sum of its activation over all tested feature set. The term *δ* = 10^−6^ in the denominator prevents division by extremely small values. The operator *max*(0, ·) discards negative activations to avoid cancellation for signed units in the sum. Thus, *r_i_* is large when unit *i* concentrates its responses on a single feature.

A unit is classified as *monosemantic* if its strength *r_i_* exceeds a threshold *τ* = 0.9 (the original study in ^33^ uses *τ* = 0.999). For a layer with *N* units, we count 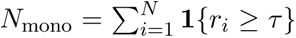. The *monosemantic score* is then defined as the number of monosemantic units per feature:

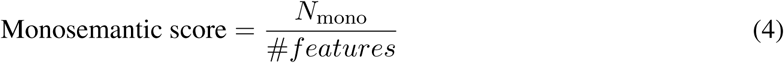

For Figure 5b where the monosemantic score was computed across all RDS types and dot densities, we used #*features* = |*T* | × |*S*| = 3 × 9 = 27. For Figure S3, where the score was computed within a given RDS type across dot densities and disparity signs, we used #*features* = |*S*| × |*D*| = 9 × 2 = 18. For the fMRI data, we used the most 250 task-sensitive voxels for each individual, the same as those used for the cross-decoding analysis. Scores were averaged across participants for fMRI data and across random seeds for DNN models.

### Statistics and reproducibility

#### Cross-decoding analysis, Fig. 2b

To ensure the statistical reliability of our cross-decoding results without relying on multivariate normality assumption that is often violated in fMRI data ^69^, we conducted a permutation test and bootstrap resampling ^70^ for the group-level analysis across participants. The statistical significance (*p*-value) of the classifier’s mean prediction accuracy for each ROI and RDS type was assessed against the chance performance baseline. The permutation test establishes a baseline for chance performance, while bootstrap resampling estimates a *p*-value for cross-decoding against a null distribution to which the true accuracy (obtained from the data) can be compared. The null distribution reflects the decoding accuracies computed when we randomized the task labels, thereby disrupting the relationship between the task labels and the voxel values ^71^. These tests require no distributional assumptions but are computationally expensive ^70^.

For the permutation test, we generated a null distribution for each aRDS-cRDS cross-decoding pair. We randomly permuted the task labels (crossed and uncrossed) within the individual’s BOLD patterns dataset for each ROI and RDS type, then performed a cross-decoding analysis using a leave-one-run-out cross-validation scheme. This process was repeated for all participants. The cross-decoding accuracies were then averaged across participants. We repeated the same procedure 10,000 times to generate a null distribution of 10,000 classification accuracies per RDS type and ROI. The baseline of chance performance level for each RDS type was defined as the 95*^th^* percentile of the corresponding null distribution.

We generated a bootstrap distribution by resampling, with replacement, individual cross-decoding performance for each ROI and RDS type. The mean of the resampled data was calculated, and this step was repeated 10,000 times. We estimated the *p*-value for each ROI and RDS type by calculating the proportion of the resulting 10,000 classification accuracies below the baseline computed in the permutation test. Following a two-tailed test and adjusting for multiple comparisons across ROIs, the confidence level alpha was set at 0.003125 [alpha = 0.05/(2 × 8 (ROIs)) = 0.003125]. A *p*-value below this threshold indicated statistical significance.

## Data and code availability

Data and codes to reproduce the results in this manuscript will be shared upon acceptance.

## Supplementary Information

### Temporal dynamics affected depth perception when processing anticorrelated features

**Figure S1:**
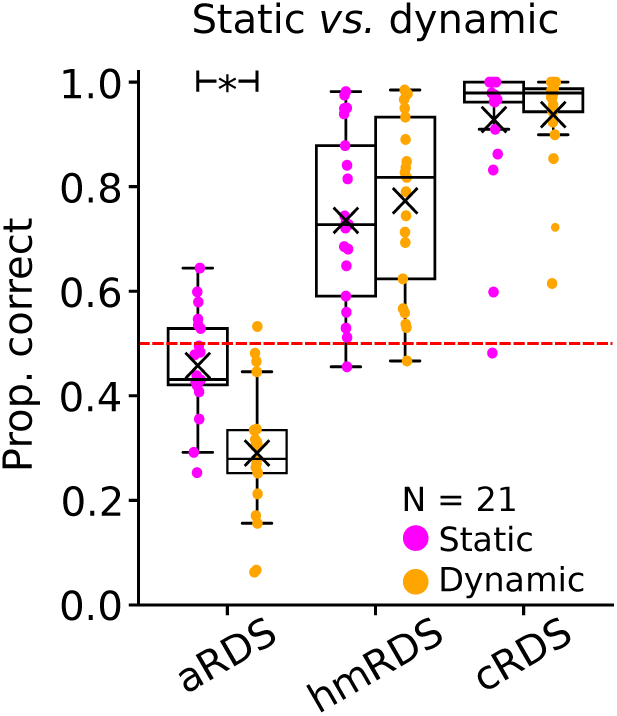
Box plots of near/far discrimination performance for aRDS, hmRDS, and cRDS under static (magenta) and dynamic (orange) conditions (*N* = 21). Each dot represents an individual participant; black crosses denote the mean; red dashed lines indicate chance (0.5). Asterisks mark a significant difference from chance (*p* = 0.0015).

The performance of depth judgment depends on many factors in visual stimuli, including the temporal dynamics. Here we conducted additional psychophysical experiments using static and dynamic anticorrelated (aRDSs), half-matched (hmRDSs), and correlated RDSs (cRDSs) in our stimulus configuration (i.e., the four circular apertures arranged in quadrants within the zero-disparity RDS background, see Fig. 1). We included cRDSs as a control condition while hmRDSs as additional parameters essential for later DNN modeling. The dot density was 25%. Each RDS type was repeated five times per run. Five participants completed 3 runs, six completed four runs, two completed five runs, eight completed six runs.

Observers reliably experienced reversed depth with dynamic aRDSs (Fig S1). Near/far discrimination for dynamic aRDSs fell significantly below chance (mean accuracy = 0.29, *p <* 0.001), indicating inversion of perceived depth (crossed-disparity was perceived far and uncrossed-disparity was perceived near, see Fig. 1b). In contrast, performance for static aRDSs did not differ significantly from chance (mean accuracy = 0.46, *p* = 0.056). A repeated-measures ANOVA examining factors of dot refresh rate (static vs. dynamic) and dot correlation revealed a significant main effect of dot refresh rate (*F*(1, 20) = 7.81, *p* = 0.012), a significant main effect of dot correlation level (*F*(2, 40) = 114.11, *p <* 0.001), and a significant interaction *F*(2, 40) = 21.18, *p <* 0.001). Post-hoc Tukey tests indicated that dynamic aRDS conditions differed significantly from their static counterparts (mean difference = 0.17, *p* = 0.0015). For cRDSs, performance was close to perfect under both static (mean accuracy = 0.93, *p <* 0.001) and dynamic (mean accuracy = 0.94, *p <* 0.001) conditions. hmRDS performance remained well above chance for both static (mean accuracy = 0.74, *p <* 0.001) and dynamic (mean accuracy = 0.77, *p <* 0.001).

These results are consistent with previous work using different stimulus configurations and different task procedures ^20,21^, and highlight the critical role of temporal dynamics in depth perception when processing anticorrelated features.

### Absolute mean difference (AMD) between human and AI psychophysics

On average across all dot densities, the DNN-Concat model showed the lowest absolute mean difference (AMD) among all models, indicating the closest overall alignment with human depth performance. For cRDSs, AMD values remained consistently low across all dot densities for all binocular interaction types, suggesting that all models captured depth representations for correlated stimuli in a manner broadly consistent with human observers.

**Figure S2:**
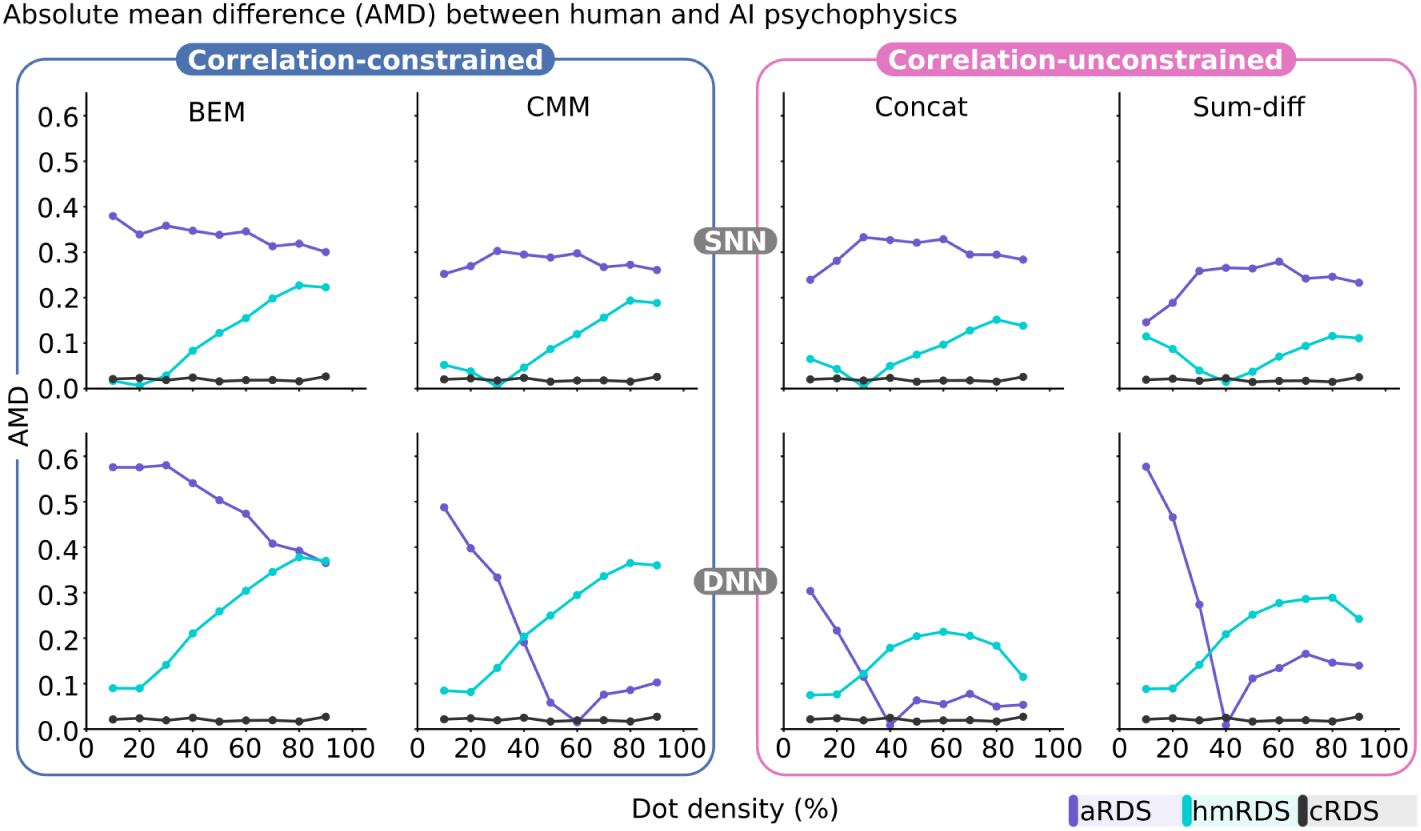
Absolute mean difference (AMD) between human and AI psychophysics.

**Table 1:** Absolute mean difference (AMD) averaged across conditions.

| Model | Mean | Lower bound | Upper bound |
| --- | --- | --- | --- |
| **SNN-BEM** | 0.158675 | 0.158632 | 0.158717 |
| **SNN-CMM** | 0.132128 | 0.132076 | 0.132180 |
| **SNN-Concat** | 0.134478 | 0.134433 | 0.134524 |
| **SNN-Sum_diff** | 0.110275 | 0.110229 | 0.110322 |
| **DNN-BEM** | 0.251526 | 0.251447 | 0.251604 |
| **DNN-CMM** | 0.149841 | 0.149774 | 0.149908 |
| **DNN-Concat** | 0.092792 | 0.092716 | 0.092868 |
| **DNN-Sum_diff** | 0.151345 | 0.151288 | 0.151403 |

For hmRDSs, the AMD for DNN-Concat increased from approximately 0.08 to 0.2 as dot density increased from 10% to 60%. Then the score decreased to around 0.1 at higher densities (60–90%). This pattern suggests that the model’s depth representations diverged from human performance at intermediate densities, where binocular matching becomes more ambiguous, but partially realigned at higher densities.

For aRDSs, the AMD decreased markedly from approximately 0.3 to near 0.01 as dot density increased from 10% to 40%, and then stabilized at around 0.05 for densities between 40% and 90%. This indicates that DNN-Concat deviated substantially from human performance at low dot densities but achieved closer alignment at moderate to high densities. Taken together, these results suggest that while DNN-Concat captures some key aspects of human depth perception, there remains important differences in processing stereo information between humans and AI.

### Voxel-level monosemanticity across brain areas

**Figure S3:**
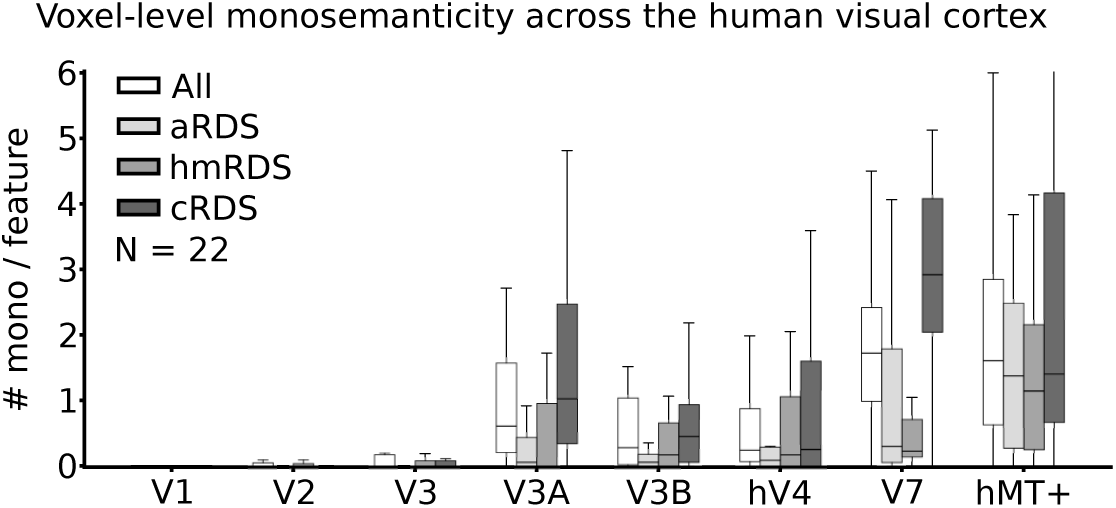
Voxel-level monosemanticity across brain areas for each RDS type.

When voxel-level monosemanticity was examined separately for each RDS type, voxel-level monosemanticity was highest for cRDS, significantly greater than aRDS (mean difference cRDS - aRDS = 0.6*, p <* 0.001), with no significant difference between hmRDS and aRDS (mean difference hmRDS - aRDS = 0.106*, p* = 0.442). This suggests that the human stereo system preferentially encodes correlated stimuli with higher specificity.

## References

[1] Julesz, B. Foundations of Cyclopean Perception (University of Chicago Press, 1971).

[2] Ohzawa, I., DeAngelis, G. C. & Freeman, R. D. Stereoscopic depth discrimination in the visual cortex: neurons ideally suited as disparity detectors. Science 249, 1037–1041 (1990).

[3] Read, J. C., Parker, A. J. & Cumming, B. G. A simple model accounts for the response of disparity-tuned V1 neurons to anticorrelated images. Vis. Neurosci. 19, 735 (2002).

[4] Read, J. Early computational processing in binocular vision and depth perception. Prog. Biophys. Mol. Biol. 87, 77–108 (2005).

[5] Doi, T., Tanabe, S. & Fujita, I. Matching and correlation computations in stereoscopic depth perception. J. Vis. 11, 1–1 (2011).

[6] Doi, T. & Fujita, I. Cross-matching: a modified cross-correlation underlying threshold energy model and match-based depth perception. Front. Comput. Neurosci. 8, 127 (2014).

[7] Henriksen, S., Cumming, B. G. & Read, J. C. A single mechanism can account for human perception of depth in mixed correlation random dot stereograms. PLoS Comput. Biol. 12, e1004906 (2016).

[8] Qian, N. Computing stereo disparity and motion with known binocular cell properties. Neural Comput. 6, 390–404 (1994).

[9] Qian, N. & Zhu, Y. Physiological computation of binocular disparity. Vision Res. 37, 1811–1827 (1997).

[10] Fujita, I. & Doi, T. Weighted parallel contributions of binocular correlation and match signals to conscious perception of depth. Philos. Trans. R. Soc. B Biol. Sci. 371, 20150257 (2016).

[11] Cumming, B. & Parker, A. Responses of primary visual cortical neurons to binocular disparity without depth perception. Nature 389, 280–283 (1997).

[12] Henriksen, S., Read, J. C. & Cumming, B. G. Neurons in striate cortex signal disparity in half-matched random-dot stereograms. J. Neurosci. 36, 8967–8976 (2016).

[13] Ohzawa, I., DeAngelis, G. C. & Freeman, R. D. Encoding of binocular disparity by complex cells in the cat’s visual cortex. J. Neurophysiol. 77, 2879–2909 (1997).

[14] Prince, S., Pointon, A., Cumming, B. & Parker, A. Quantitative analysis of the responses of V1 neurons to horizontal disparity in dynamic random-dot stereograms. J. Neurophysiol. 87, 191–208 (2002).

[15] Prince, S., Cumming, B. & Parker, A. Range and mechanism of encoding of horizontal disparity in macaque V1. J. Neurophysiol. 87, 209–221 (2002).

[16] Tanabe, S., Umeda, K. & Fujita, I. Rejection of false matches for binocular correspondence in macaque visual cortical area V4. J. Neurosci. 24, 8170–8180 (2004).

[17] Elhage, N., et al. Toy models of superposition. arXiv preprint arXiv:2209.10652 (2022).

[18] Read, J. C. & Cumming, B. G. Sensors for impossible stimuli may solve the stereo correspondence problem. Nat. Neurosci. 10, 1322–1328 (2007).

[19] Goncalves, N. R. & Welchman, A. E. “what not” detectors help the brain see in depth. Curr. Biol. 27, 1403–1412 (2017).

[20] Doi, T., Takano, M. & Fujita, I. Temporal channels and disparity representations in stereoscopic depth perception. J. Vis. 13, 26–26 (2013).

[21] Zhaoping, L. Testing the top-down feedback in the central visual field using the reversed depth illusion. iScience 28 (2025).

[22] Wundari, B. G., Fujita, I. & Ban, H. Human and artificial visual systems share a computational principle for transforming binocular disparity into depth representation. *Commun*. Biol. 8, 1042 (2025).

[23] Janssen, P., Vogels, R., Liu, Y. & Orban, G. A. At least at the level of inferior temporal cortex, the stereo correspondence problem is solved. Neuron 37, 693–701 (2003).

[24] Takemura, A., Inoue, Y., Kawano, K., Quaia, C. & Miles, F. Single-unit activity in cortical area MST associated with disparity-vergence eye movements: evidence for population coding. J. Neurophysiol. 85, 2245–2266 (2001).

[25] Krug, K., Cumming, B. G. & Parker, A. J. Comparing perceptual signals of single V5/MT neurons in two binocular depth tasks. J. Neurophysiol. 92, 1586–1596 (2004).

[26] Yoshioka, T. W., Doi, T., Abdolrahmani, M. & Fujita, I. Specialized contributions of mid-tier stages of dorsal and ventral pathways to stereoscopic processing in macaque. eLife 10, e58749 (2021).

[27] Kriegeskorte, N., Goebel, R. & Bandettini, P. Information-based functional brain mapping. Proc. Natl Acad. Sci. USA 103, 3863–3868 (2006).

[28] Chirimuuta, M. Explanation in computational neuroscience: Causal and non-causal. Br. J. Philos. Sci. (2018).

[29] Cohn, T. & Lasley, D. Binocular vision: Two possible central interactions between signals from two eyes. Science 192, 561–563 (1976).

[30] May, K. A. & Zhaoping, L. Efficient coding theory predicts a tilt aftereffect from viewing untilted patterns. Curr. Biol. 26, 1571–1576 (2016).

[31] Mayer, N. et al. A large dataset to train convolutional networks for disparity, optical flow, and Scene Flow estimation. Proc. IEEE Conf. Comput. Vis. Pattern Recognit. 4040–4048 (2016). ArXiv:1512.02134.

[32] Bengio, Y., Courville, A. & Vincent, P. Representation learning: A review and new perspectives. IEEE Trans. Pattern Anal. Mach. Intell. 35, 1798–1828 (2013).

[33] Jermyn, A. S., Schiefer, N. & Hubinger, E. Engineering monosemanticity in toy models. arXiv preprint arXiv:2211.09169 (2022).

[34] Haefner, R. M. & Cumming, B. G. Adaptation to natural binocular disparities in primate V1 explained by a generalized energy model. Neuron 57, 147–158 (2008).

[35] DeAngelis, G. C., Ohzawa, I. & Freeman, R. D. Depth is encoded in the visual cortex by a specialized receptive field structure. Nature 352, 156 (1991).

[36] Saxe, A. M., McClelland, J. L. & Ganguli, S. Exact solutions to the nonlinear dynamics of learning in deep linear neural networks. arXiv preprint arXiv:1312.6120 (2013).

[37] Barak, O., Rigotti, M. & Fusi, S. The sparseness of mixed selectivity neurons controls the generalization–discrimination trade-off. J. Neurosci. 33, 3844–3856 (2013).

[38] Spanne, A. & Jörntell, H. Questioning the role of sparse coding in the brain. Trends Neurosci. 38, 417–427 (2015).

[39] Johnston, W. J., Palmer, S. E. & Freedman, D. J. Nonlinear mixed selectivity supports reliable neural computation. PLoS Comput. Biol. 16, e1007544 (2020).

[40] Hubel, D. H. & Wiesel, T. N. Receptive fields and functional architecture of monkey striate cortex. J. Physiol. 195, 215–243 (1968).

[41] Dougherty, K., Cox, M. A., Westerberg, J. A. & Maier, A. Binocular modulation of monocular V1 neurons. Curr. Biol. 29, 381–391 (2019).

[42] Zhang, S.-H., Zhao, X.-N., Jiang, D.-Q., Tang, S.-M. & Yu, C. Ocular dominance-dependent binocular combination of monocular neuronal responses in macaque V1. eLife 13, RP92839 (2024).

[43] Tang, S. et al. Complex pattern selectivity in macaque primary visual cortex revealed by large-scale two-photon imaging. Curr. Biol. 28, 38–48 (2018).

[44] Brincat, S. L., Siegel, M., von Nicolai, C. & Miller, E. K. Gradual progression from sensory to task-related processing in cerebral cortex. Proc. Natl Acad. Sci. USA 115, E7202–E7211 (2018).

[45] Kriegeskorte, N., Cusack, R. & Bandettini, P. How does an fMRI voxel sample the neuronal activity pattern: compact-kernel or complex spatiotemporal filter? NeuroImage 49, 1965–1976 (2010).

[46] Goncalves, N. R. et al. 7 tesla fmri reveals systematic functional organization for binocular disparity in dorsal visual cortex. J. Neurosci. 35, 3056–3072 (2015).

[47] Alvarez, I., Mancari, A., Ip, I. B., Parker, A. J. & Bridge, H. Characterizing human disparity tuning properties using population receptive field mapping. J. Neurosci. 45 (2025).

[48] Parker, A. J., et al. Receptive fields from single-neuron recording and MRI reveal similar information coding for binocular depth. Proc. Natl Acad. Sci. USA 122, e2409893122 (2025).

[49] Kleiner, M., Brainard, D. & Pelli, D. What’s new in Psychtoolbox-3? Perception 36, 14 (2007). ECVP Abstract Supplement.

[50] Ban, H. & Yamamoto, H. A non–device-specific approach to display characterization based on linear, nonlinear, and hybrid search algorithms. J. Vis. 13, 20–20 (2013).

[51] Popple, A. V., Smallman, H. S. & Findlay, J. M. The area of spatial integration for initial horizontal disparity vergence. Vision Res. 38, 319–326 (1998).

[52] Tanabe, S., Yasuoka, S. & Fujita, I. Disparity-energy signals in perceived stereoscopic depth. J. Vis. 8, 22–22 (2008).

[53] Sereno, M. I. et al. Borders of multiple visual areas in humans revealed by functional magnetic resonance imaging. Science 268, 889–893 (1995).

[54] DeYoe, E. A., et al. Mapping striate and extrastriate visual areas in human cerebral cortex. Proc. Natl Acad. Sci. USA 93, 2382–2386 (1996).

[55] Dumoulin, S. O. & Wandell, B. A. Population receptive field estimates in human visual cortex. NeuroImage 39, 647–660 (2008).

[56] Yamamoto, H. et al. Large-and small-scale functional organization of visual field representation in the human visual cortex. Visual cortex: New research 195–226 (2008).

[57] Huk, A. C., Dougherty, R. F. & Heeger, D. J. Retinotopy and functional subdivision of human areas MT and MST. J. Neurosci. 22, 7195–7205 (2002).

[58] Ban, H., Preston, T. J., Meeson, A. & Welchman, A. E. The integration of motion and disparity cues to depth in dorsal visual cortex. Nat. Neurosci. 15, 636 (2012).

[59] Kendall, A. et al. End-to-end learning of geometry and context for deep stereo regression. Proc. IEEE Int. Conf. Comput. Vis. 66–75 (2017).

[60] Rigotti, M. et al. The importance of mixed selectivity in complex cognitive tasks. Nature 497, 585–590 (2013).

[61] Fusi, S., Miller, E. K. & Rigotti, M. Why neurons mix: high dimensionality for higher cognition. Curr. Opin. Neurobiol. 37, 66–74 (2016).

[62] Tye, K. M. et al. Mixed selectivity: Cellular computations for complexity. Neuron 112, 2289–2303 (2024).

[63] Kira, S., Safaai, H., Morcos, A. S., Panzeri, S. & Harvey, C. D. A distributed and efficient population code of mixed selectivity neurons for flexible navigation decisions. Nat. Commun. 14, 2121 (2023).

[64] Yiling, Y., Klon-Lipok, J. & Singer, W. Joint encoding of stimulus and decision in monkey primary visual cortex. Cereb. Cortex 34, bhad420 (2024).

[65] Franke, K. et al. Dual-feature selectivity enables bidirectional coding in visual cortical neurons. eLife 15, RP109861 (2026).

[66] Zhang, C. Y. et al. Partially mixed selectivity in human posterior parietal association cortex. Neuron 95, 697–708 (2017).

[67] Dang, W., Li, S., Pu, S., Qi, X.-L. & Constantinidis, C. More prominent nonlinear mixed selectivity in the dorsolateral prefrontal than posterior parietal cortex. eNeuro 9 (2022).

[68] Scherlis, A., Sachan, K., Jermyn, A. S., Benton, J. & Shlegeris, B. Polysemanticity and capacity in neural networks. arXiv preprint arXiv:2210.01892 (2022).

[69] Kriegeskorte, N. Pattern-information analysis: from stimulus decoding to computational-model testing. NeuroImage 56, 411–421 (2011).

[70] Nichols, T. E. & Holmes, A. P. Nonparametric permutation tests for functional neuroimaging: a primer with examples. Hum. Brain Mapp. 15, 1–25 (2002).

[71] Etzel, J. A. MVPA permutation schemes: permutation testing for the group level. Proc. Int. Workshop Pattern Recognit. Neuroimaging 65–68 (2015).

